# KRAS-mediated upregulation of CIP2A promotes suppression of PP2A-B56α to initiate pancreatic cancer development

**DOI:** 10.1101/2023.07.01.547283

**Authors:** Samantha L. Tinsley, Rebecca A. Shelley, Gaganpreet K. Mall, Ella Rose D. Chianis, Alisha Dhiman, Garima Baral, Harish Kothandaraman, Mary C. Thoma, Colin J. Daniel, Nadia Atallah Lanman, Marina Pasca di Magliano, Goutham Narla, Luis Solorio, Emily C. Dykhuizen, Rosalie C. Sears, Brittany L. Allen-Petersen

## Abstract

Oncogenic mutations in KRAS are present in approximately 95% of patients diagnosed with pancreatic ductal adenocarcinoma (PDAC) and are considered the initiating event of pancreatic intraepithelial neoplasia (PanIN) precursor lesions. While it is well established that KRAS mutations drive the activation of oncogenic kinase cascades during pancreatic oncogenesis, the effects of oncogenic KRAS signaling on regulation of phosphatases during this process is not fully appreciated. Protein Phosphatase 2A (PP2A) has been implicated in suppressing KRAS-driven cellular transformation. However, low PP2A activity is observed in PDAC cells compared to non-transformed cells, suggesting that suppression of PP2A activity is an important step in the overall development of PDAC. In the current study, we demonstrate that KRAS^G12D^ induces the expression of both an endogenous inhibitor of PP2A activity, Cancerous Inhibitor of PP2A (CIP2A), and the PP2A substrate, c-MYC. Consistent with these findings, KRAS^G12D^ sequestered the specific PP2A subunit responsible for c-MYC degradation, B56α, away from the active PP2A holoenzyme in a CIP2A-dependent manner. During PDAC initiation *in vivo*, knockout of B56α promoted KRAS^G12D^ tumorigenesis by accelerating acinar-to-ductal metaplasia (ADM) and the formation of PanIN lesions. The process of ADM was attenuated *ex vivo* in response to pharmacological re-activation of PP2A utilizing direct small molecule activators of PP2A (SMAPs). Together, our results suggest that suppression of PP2A-B56α through KRAS signaling can promote the MYC-driven initiation of pancreatic tumorigenesis.

## INTRODUCTION

The development of mutations in protooncogene Kirsten Rat Sarcoma viral oncogene homolog (KRAS) is known to be a driver of many common cancers, such as lung cancer, colorectal cancer, and pancreatic cancer (1). Pancreatic ductal adenocarcinoma (PDAC) has the lowest five-year survival rate (12%) of all major cancers (2). Within PDAC tumors, oncogenic KRAS mutations occur in approximately 95% of all patients, underscoring the importance of this signaling pathway to tumor development, progression, and patient outcomes (3). Data generated from genetic mouse models have established that the expression of mutant KRAS in the pancreas drives the initiation of pancreatic cancer and helps to maintain malignant tumor growth (4,5). Acinar-to-ductal metaplasia (ADM) is a reversible process through which pancreatic acinar cells undergo a transient transdifferentiation process to a duct-like state in response to exogenous stressors, such as wound infliction or inflammation (6). However, in the presence of oncogenic KRAS, this process becomes permanent and leads to the formation of pancreatic intraepithelial neoplasia (PanIN) precursor lesions (6). Historically, KRAS mutations alone have not been sufficient to transform normal cells into malignant cells and instead often cause oncogene-induced senescence (4,7). However, inhibition of Protein Phosphatase 2A (PP2A) combined with oncogenic KRAS mutations is adequate for malignant transformation, implicating PP2A as a critical negative regulator of KRAS signaling (8).

PP2A is a heterotrimeric holoenzyme composed of three individual subunits that together comprise almost 90 unique complexes: the (A) scaffolding subunit, the (B) substrate specifying regulatory subunit, and the (C) catalytic subunit (9). The B subunits are separated into four major families: B (B55), B’ (B56), B’’, and B’’’(Striatins). The impact of PP2A activity on oncogenic signaling heavily depends on which B subunit is incorporated into the complex and is highly context specific. Using a targeted shRNA screen, Sablina et al. determined that loss of B56α could transform human embryonic kidney cells expressing HRAS^G12V^, implicating the PP2A-B56α complex as a tumor suppressor (8). While the B56α subunit dephosphorylates many downstream KRAS signaling effectors, including MEK, ERK, AKT, and the oncoprotein c-MYC (MYC), the functional role of this subunit during PDAC progression is relatively unknown (10–13). Loss of PP2A is associated with cellular transformation; however genetic deletion or inactivating mutations of PP2A subunits in pancreatic cancer is extremely low (9). Conversely, we and others have shown that endogenous inhibitors of PP2A, such as Cancerous Inhibitor of PP2A (CIP2A), are commonly overexpressed in PDAC and contribute to tumor growth (14). Furthermore, PDAC tumors commonly display aberrantly high levels of active phosphorylated MYC without associated genetic amplification of the gene (14). Therefore, we hypothesize that PP2A functions as a critical posttranslational gatekeeper to maintain cellular homeostasis. In the current study, we aim to understand how the suppression of PP2A intersects with oncogenic KRAS to promote the initiation of pancreatic tumorigenesis.

Here, we demonstrate that induction of KRAS^G12D^ in human pancreatic epithelial cells increases the expression of both CIP2A and the direct B56α substrate, MYC. The increased CIP2A expression is associated with decreased incorporation of B56α into the PP2A holoenzyme and is rescued with CIP2A knockdown. To determine the functional consequence of B56α depletion *in vivo*, we generated a novel KRAS^G12D^-driven mouse model of PDAC with hypomorphic loss of B56α (B56α^hm/hm^ (15) in the pancreas. Loss of B56α significantly accelerated KRAS^G12D^-driven ADM and PanIN formation. Finally, treatment of ADM cultures *ex vivo* with the small molecule activator of PP2A, DT061, prevented the transdifferentiation of acinar cell to a ductal-like state. Together, these findings implicate the inhibition of PP2A-B56α as a critical early event in PDAC initiation and highlight the understudied potential of protein phosphatases as actionable therapeutic targets.

## RESULTS

### Induction of KRAS^G12D^ promotes CIP2A-mediated suppression of PP2A-B56α in pancreatic epithelial cells

Previous studies have suggested that CIP2A and KRAS may function synergistically and display a significant overlap in downstream regulated pathways, suggesting intersecting signaling pathway regulation (16–18). Additionally, in KRAS mutant PDAC patients, low B56α expression correlates with poor prognosis (**Fig S1A**). Therefore, to determine the acute effects of oncogenic KRAS on CIP2A expression, a non-transformed human pancreatic ductal epithelial cell line (HPDE) with tet-inducible KRAS^G12D^ (HPDE-iKRAS) was treated with doxycycline (dox) in a time course experiment. This system allowed for potent induction of KRAS^G12D^ expression (**Fig S1B**) and acute activation of the downstream mitogen-activated protein kinase (MAPK) pathway, with peak phospho-ERK (pERK, p-p42/44) levels seen at 8H post-dox treatment. Induction of KRAS^G12D^ resulted in a time-dependent increase in CIP2A expression, with significant upregulation seen by 8-12hr (**Fig 1A, B, D**). PP2A-B56α provides critical posttranslational regulation of MYC by dephosphorylating MYC at Serine62 (pS62 MYC) and driving MYC proteasomal degradation (19,20). Additionally, this specific PP2A heterotrimer is negatively regulated by CIP2A (21). Consistent with increased expression of CIP2A and PP2A inhibition, KRAS^G12D^ induction significantly increased MYC protein expression (**Fig 1A, C**) in a manner that mirrored CIP2A kinetics. In contrast, there was no change in *MYC* mRNA levels (**Fig S1B**), supporting a posttranslational regulatory event.

**Figure 1:**
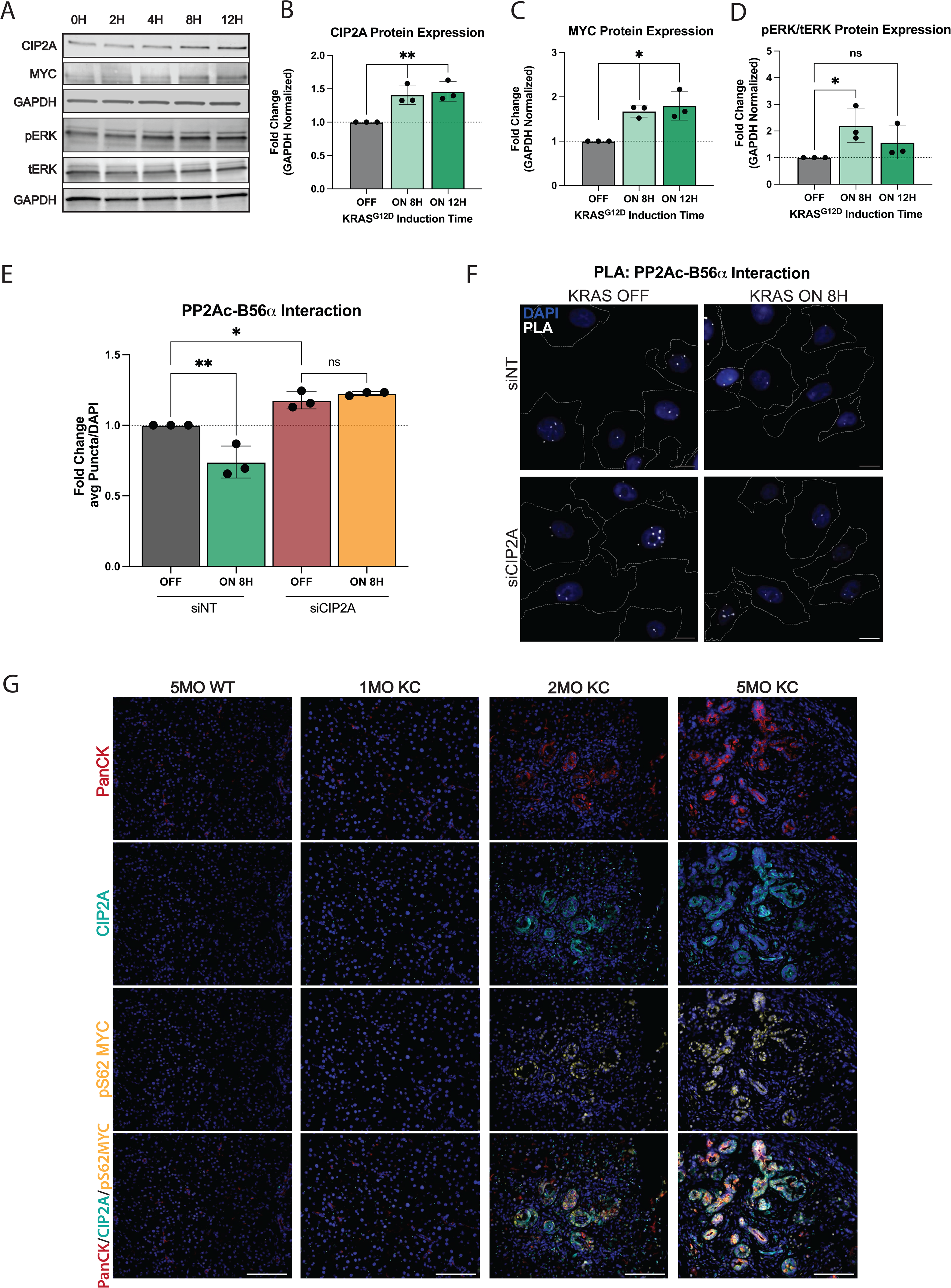
Oncogenic KRAS signaling promotes suppression of the PP2A-B56a complex. **(A)** Representative western blot of human pancreatic ductal epithelial (HPDE) cells with tet-inducible KRAS^G12D^ (HPDE-iKRAS) treated with 50ng/mL doxycycline for 0-12H in a time course. **(B-D)** Quantification of CIP2A, MYC, and pERK/tERK protein expression at 0, 8, 12H post-KRAS^G12D^ induction in HPDE-iKRAS cells. **(E)** Quantification of average puncta per cell of PLA between B56α and PP2Ac after 0H (OFF) or 8H (ON) of KRASG12D induction and treated with siNT siRNA or siRNA to CIP2A. Quantification is normalized to the average signal from the siNT OFF condition. **(F)** Representative images of PLA quantified in E. Scale bars represent 10μm, white dashed lines represent edges of cells as identified through actin staining. **(G)** Representative images of three independent biological replicate stains of WT, 1MO, 2MO, and 5MO KC pancreatic tissue (left to right). IF stains indicate PanCK, CIP2A, pS62 MYC, and MERGE of all stains (top to bottom). Scale bar represents 100μm. All experiments were repeated as three independent biological replicates and statistical analysis was performed using one-way ANOVA. \**p*<0.05 \*\**p*<0.01 n.s. = not significant

To identify the impact of KRAS^G12D^ on the ability of cells to form an active PP2A-B56α holoenzyme, HPDE-iKRAS cells were treated with dox for 8H and the interaction between the catalytic subunit of PP2A (PP2Ac) and B56α was determined using a proximity ligation assay (PLA). Dox treatment significantly reduced the association of B56α with PP2Ac, as seen by decreased PLA puncta, supporting the functional inhibition of PP2A-B56α in response to KRAS induction (**Fig 1E-F**). This reduction in PP2Ac-B56α was dependent on CIP2A, as knockdown of CIP2A with siRNA (siCIP2A) significantly increased the number of PP2Ac-B56α complexes in (-) dox conditions and prevented the KRAS^G12D^-induced loss of PP2Ac-B56α association in (+) dox conditions (**Figure 1E-F, S1E**). Furthermore, transient knock down of CIP2A (siCIP2A) reduced clonogenic colony growth in HPDE-iKRAS cells compared to non-targeting siRNA (siNT) treated cells (**Fig S1C-D**), indicating that CIP2A expression supports the self-renewal capacity of these cells.

To identify whether KRAS^G12D^ expression correlates with lower PP2A function/activity during pancreatic cancer disease progression, we utilized the *PDX1-Cre; LSL-KRAS^G12D^* (KC (4) pancreatic cancer mouse model, which conditionally expresses KRAS^G12D^ in all cells of the pancreas. These mice develop robust ADM and PanIN lesions that are histologically similar to human disease. KC pancreatic tissues from mice at 1-, 2-, and 5-months of age were stained for CIP2A (PP2A inhibitor) and pS62 MYC (PP2A target), and pan-Cytokeratin (epithelial populations) (**Fig 1G**). Compared to normal pancreata from WT mice, there was a significant increase in both CIP2A and pS62 MYC protein expression within precursor PanIN lesions. Furthermore, this expression increased across time as PanINs progressed (**Fig 1G**). Together, these findings indicate that the functional inhibition of B56α downstream of KRAS^G12D^ is an early event in PDAC development.

### Loss of KRAS^G12D^ alleviates CIP2A-mediated suppression of B56α

To identify whether KRAS^G12D^ signaling is necessary to maintain CIP2A-mediated suppression of PP2A-B56α, we utilized a KRAS-addicted malignant cell line (iKRAS3) isolated from the tet-inducible KRAS^G12D^ PDAC mouse model (iKras*; *p48*-Cre;R26-rtTa-IRES-EGFP;TetO- *Kras^G12D^*(5)) In this system, Kras^G12D^ expression is maintained by continuously culturing cells in dox-containing media and the removal of dox results in a rapid loss of Kras^G12D^ expression. iKras3 cells were cultured with dox and then harvested 0, 24, 48, and 72 hours (H) post-dox withdrawal. Consistent with dox-inducible Kras^G12D^ in Figure 1, pErk levels returned to baseline within 24H after the loss of transgenic Kras^G12D^ expression (tKras^G12D^) (**Fig 2A-C**). Conversely, B56α expression increased by 24H post-dox and remained elevated for the rest of the time course. Inhibition of the MAPK pathway is known to attenuate MYC activity and stability (22). However, despite pEKR levels being low, MYC levels remained high 24H post-dox withdrawal, and then declined over time. The sustained MYC signaling at this timepoint correlates with a transient increase in CIP2A expression at 24H that then rapidly dropped off, with a 60% inhibition of MYC occurring by 72H (**Fig 2A-B**). While the loss of CIP2A at 72H could be due to decreased transcription, *Myc* mRNA remained unchanged after 72H (**Fig 2C**).

**Figure 2:**
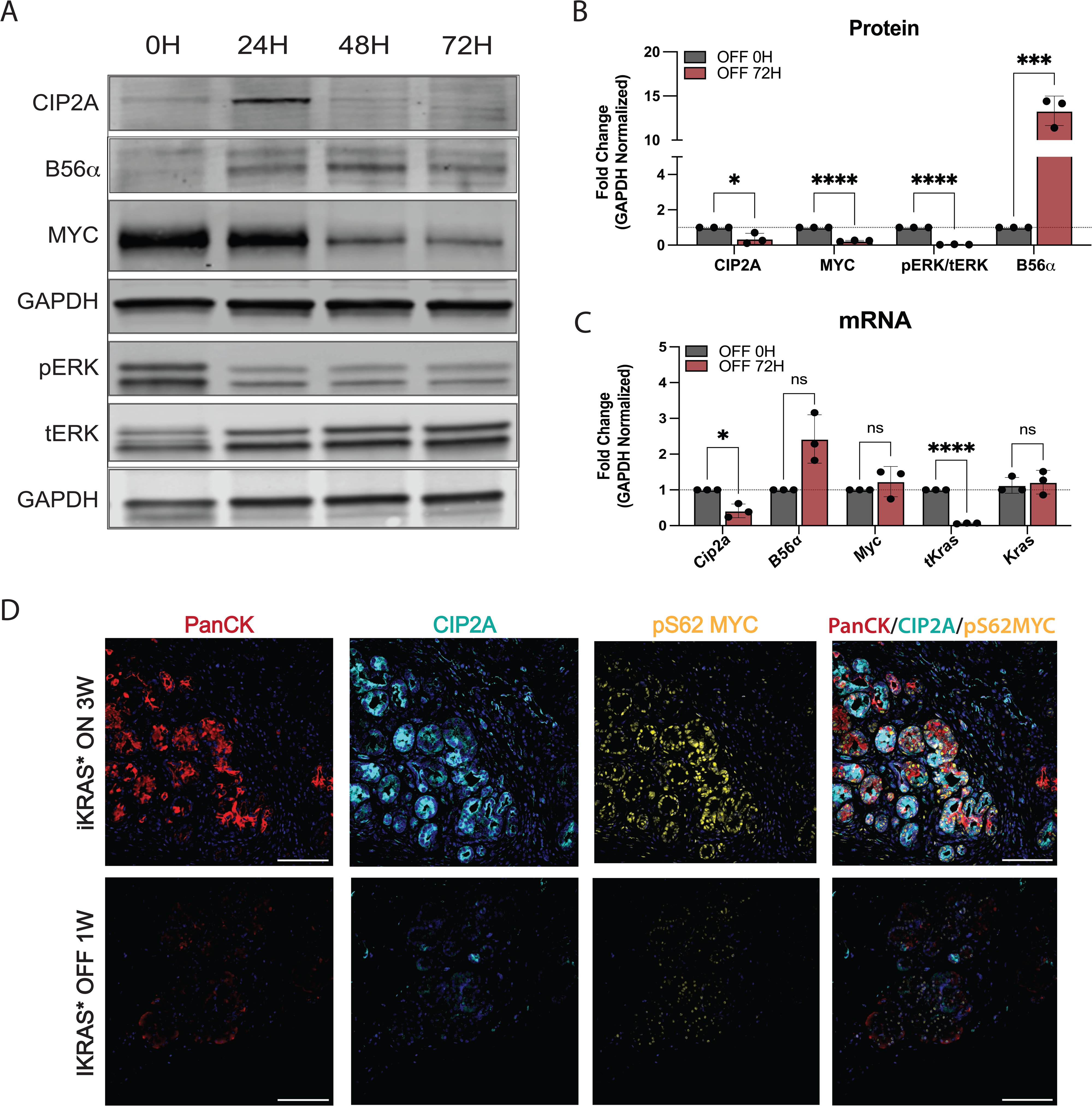
Loss of oncogenic KRAS signaling promotes activation of PP2A-B56a. **(A)** Representative western blot of protein signaling in tet-inducible KRAS^G12D^ after 0-72H after dox removal (loss of KRAS^G12D^) in iKRAS3 cells. **(B)** Quantification of three independent replicates of western blot represented in A at 0H and 72H without KRAS^G12D^ signaling. **(C)** Matching mRNA isolated from iKRAS3 cells with loss of KRAS^G12D^ signaling for 72H. Quantification includes three independent biological replicates. **(E)** Representative images of pancreatic tissue from iKRAS* mouse models after 3W of dox-induced KRAS^G12D^ signaling (top) or after 1W of dox withdrawal (bottom). Immunofluorescent staining of PanCK, CIP2A, pS62 MYC, and all stains merged (left to right). Statistical analysis was performed using one-way ANOVA. All data are representative of 3-4 biological replicates. \**p*<0.05 \*\*\**p*<0.001 \*\*\*\**p*<0.0001 n.s. = not significant. Scale bar represents 100μm.

To determine if loss of KRAS^G12D^ results in a similar decrease in CIP2A and MYC protein expression *in vivo*, we induced Kras^G12D^ for three weeks in the pancreas of adult iKras* mice and then harvested pancreatic tissue one week after dox withdrawal (**Fig 2D**). Loss of Kras^G12D^ in this model leads to a dramatic remodeling of the pancreas, with decreased PanINs and a re-emergence of acinar tissue (5). Compared to +dox conditions, loss of Kras^G12D^ expression dramatically reduced both CIP2A and active MYC (pS62 MYC) protein expression. Together, these results demonstrate that oncogenic KRAS drives and maintains CIP2A expression, suggesting that suppression of PP2A-B56α is likely important for initiating and promoting PDAC progression.

### Overexpression of PP2A-B56α abrogates KRAS^G12D^ tumorigenic phenotypes

Given that induction of KRAS signaling leads to the suppression of PP2A-B56α activation, we next sought determine if genetic overexpression of the PP2A B56α subunit would be sufficient to abrogate oncogenic KRAS phenotypes. Therefore, we overexpressed an HA-tagged B56α (HA-B56α) in malignant iKRAS3 cells (iKRAS3 B56αOE) and compared overall tumorigenic properties relative to empty vector controls (iKRAS3 EV) (23). First, we evaluated the effects of B56α overexpression on both anchorage-independent growth and self-renewal capabilities *in vitro* through soft agar (**Fig 3A-B**) and clonogenic assays (**Fig 3C-D**), respectively. We found that overexpression of PP2A-B56α significantly abrogated the ability of iKRAS3 to grow in both soft agar and clonogenic assays, indicating that suppression of PP2A-B56α expression is important for maintaining oncogenic KRAS^G12D^ phenotypes.

**Figure 3:**
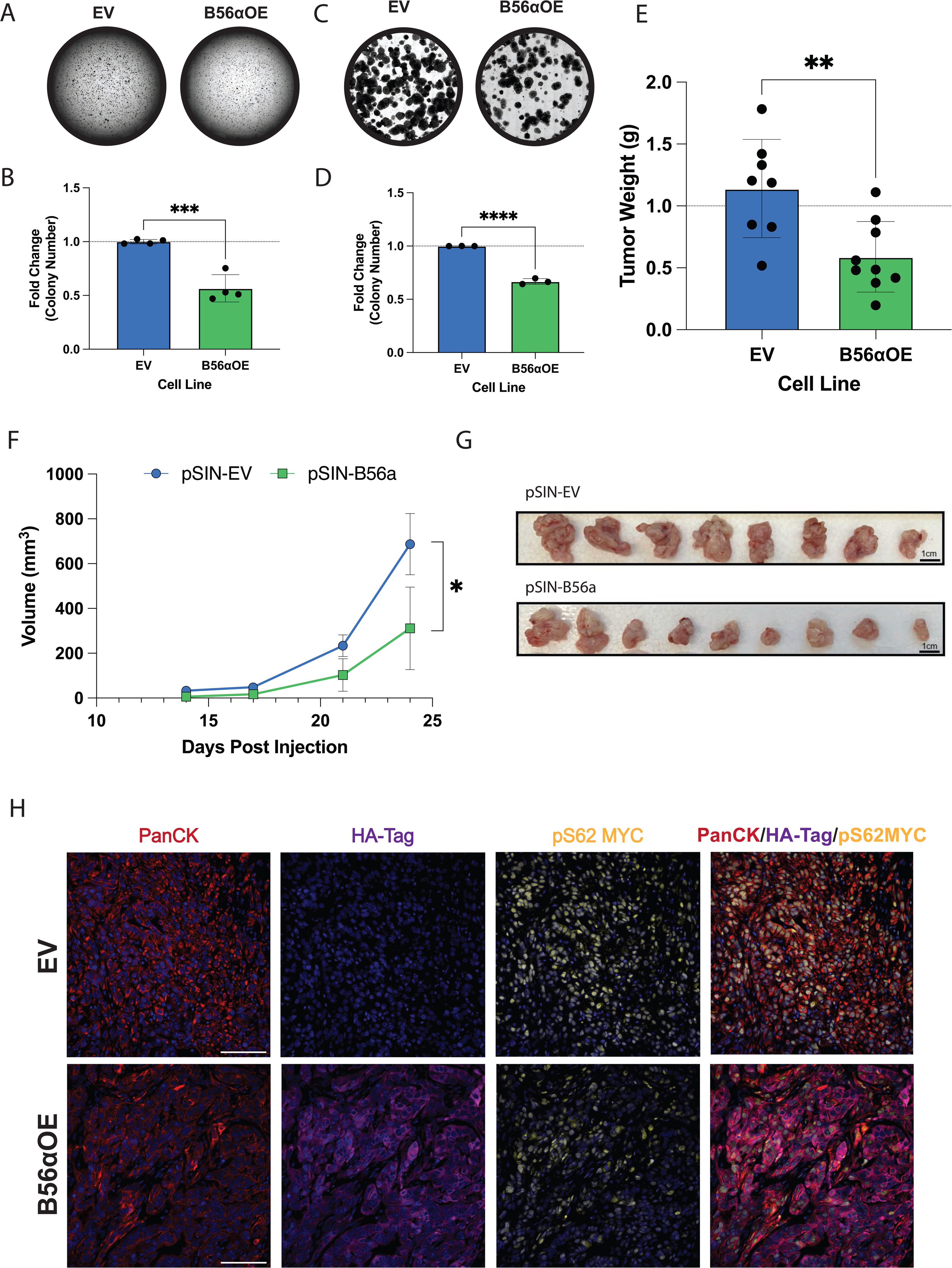
Overexpression of B56a abrogates oncogenic phenotypes. **(A)** Soft agar assay of stable iKRAS3 cell line with empty vector control or overexpression of B56α, stained with crystal violet and scanned after two weeks of growth. **(B)** Quantification of four biological replicates of the soft agar assay represented as fold change relative to the number of colonies formed in the iKRAS3 empty vector control. **(C)** Clonogenic assay of iKRAS3 cells with empty vector control or overexpression of B56α stained with crystal violet after six days. **(D)** Quantification of three replicates of clonogenic assay, measured by fold change in the percent area covered by cells. Quantified using FIJI. **(E)** Endpoint tumor weight in grams of each condition. (EV *n* = 8, B56α OE *n* = 9). **(F)** Growth curve of orthotopic pancreatic allograft of iKRAS 3 EV or B56αOE cell lines in NRG mice, volume of tumor in mm^3^. **(G)** Picture of orthotopic tumors at endpoint displayed from largest (right) to smallest (left) in each condition (top: EV, bottom: B56αOE). **(H)** Representative images of immunofluorescent staining from EV (top) or B56αOE (bottom) tumors. Stains in images represent PanCK, HA-tag, pS62 MYC, and merged images (left to right). All experiments were performed at least three times independently and statistical analysis was done using student’s t-test. \**p*<0.05 \*\**p*<0.01 \*\*\**p*<0.001 \*\*\*\**p*<0.0001 Scale bar represents 100μm.

To test if B56α expression suppresses KRAS^G12D^ *in vivo* tumor growth, iKRAS3 EV or B56αOE cell were orthotopically injected into the pancreas of *NOD.Cg-Rag1^tm1Mom^ Il2rg^tm1Wjl^/SzJ* (NRG) mice and tumor growth was measured weekly by high-frequency ultrasound imaging. Overexpression of B56α significantly hindered *in vivo* tumor growth kinetics, resulting in a 50% reduction in tumor weight at endpoint (**Fig 3E-G**). Immunofluorescent staining of isolated tumor tissue in both the EV and B56αOE conditions showed that areas of HA-B56α expression correlate with lower levels of pS62 MYC (**Fig 3H**), demonstrating that increased B56α expression antagonizes MYC protein expression downstream of Kras^G12D^, leading to abrogated tumor growth.

### Genetic loss of PP2A-B56α accelerates KRAS^G12D^-driven acinar-to-ductal metaplasia *ex vivo*

Acinar-to-ductal metaplasia (ADM) represents one of the earliest steps in PDAC progression and is irreversibly driven by oncogenic KRAS (4). *Ex vivo* ADM assays have proven to be an excellent system to study ADM kinetics and the signaling mechanisms that contribute to this process (24). ADM assays are performed by isolating pancreatic acinar cells, either from human or mouse tissues, and embedding them into a collagen I matrix. Using this technique, previous reports have shown that in response to stimuli (e.g. TGF-a, RANTES, etc.) or expression of KRAS^G12D^, pancreatic acinar cells transdifferentiate to a duct-like state and exhibit dramatic morphological and transcriptional changes. To determine the impact of B56α loss on KRAS^G12D^- driven ADM, we isolated acinar cells from 5-6 week old WT, B56α hypomorph mice (B56α^hm/hm^ (15)), *PDX1-Cre;LSL-KRAS^G12D^* (KC (4)), or *PDX1-Cre;LSL-Kras^G12D^;B56α^hm/hm^* (KCB^hm/hm^) mice and implanted the acinar clusters into a collagen I matrix per established protocols (24). The acinar clusters were imaged every six hours for three days and the number of acinar and ductal structures in each condition were quantified at 24, 48, and 72H post plating. Consistent with previous reports, without expression of *Kras^G12D^*, acinar cells derived from WT tissues were unable to transdifferentiate and retained their acinar identity throughout the 72-hour time course (25). Similar results were seen in the B56α^hm/hm^ cultures, indicating that the genetic loss of B56α is not sufficient to drive the ADM process (**Fig 4A-B)**. In contrast, loss of B56α significantly accelerated Kras^G12D^-driven ADM, with duct-like structures emerging almost 24 hours earlier in KCB^hm/hm^ cells compared to KC (**Fig 4A-B, S2A**). However, both KC and KCB^hm/hm^ conditions had nearly completed the differentiation process after 72H of implantation. To validate the transdifferentiation from an acinar to a ductal cell identity, cells were isolated out of the collagen matrix at 24 and 48H and mRNA gene expression was quantified over time. There was a significant increase in the ductal marker, cytokeratin 19 (*Ck19)* (**Fig 4C**), and the ductal transcription factor, sry-box transcription factor 9 (*Sox9)* (**Fig S2B**) over time as ductal structures emerged. Consistent with the accelerated ADM phenotype, there was a significant upregulation in *Ck19* in KCB^hm/hm^ cells compared to KC alone. As structures differentiate from acinar to ductal, *Cip2a* and *Myc* mRNA are also upregulated (**Fig S2C, D**), suggesting that the dysregulation of the CIP2A-PP2A-MYC signaling axis contributes to the transdifferentiation to a duct-like cell fate.

**Figure 4:**
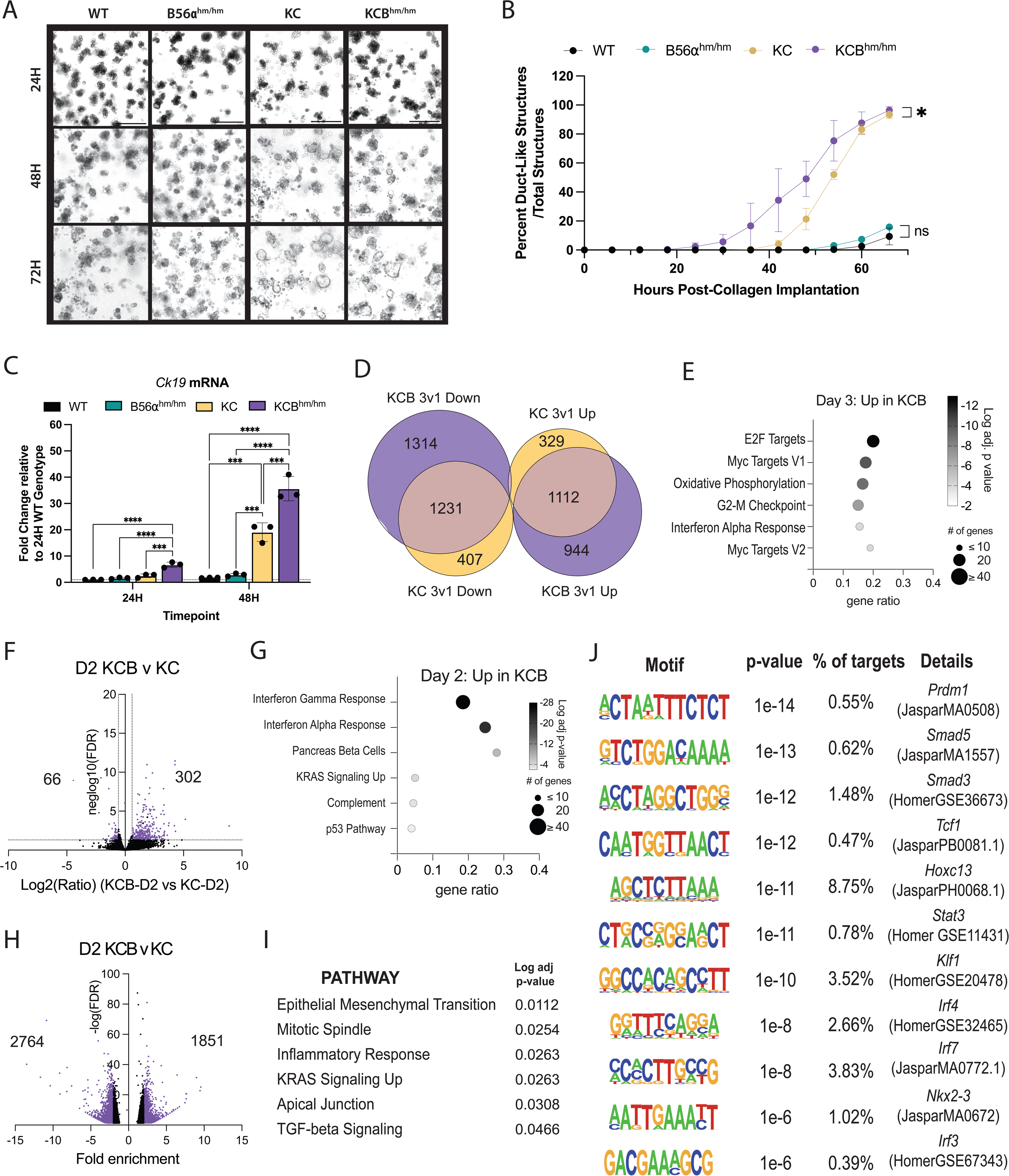
Loss of B56a promotes acceleration of acinar-to-ductal metaplasia. **(A)** Representative images of acinar cells isolated from WT, B56a^hm/hm^, KC, and KCB^hm/hm^ genetic mouse models cultured in collagen for 24, 48, and 72H. **(B)** Time course of acinar- to-ductal transdifferentiation kinetics in each condition over three days, quantified every 6H. Time course includes data from one representative biological replicate. **(C)** mRNA expression of ductal cell marker, *ck19*, in each genotype at 24 and 48H timepoints. **(D)** Venn Diagram of differentially regulated genes (DEGs) increased (UP) or decreased (DOWN) from Day 1 versus Day 3 comparisons of each genotype. Overlapping DEGs represented where Venn Diagram overlaps. **(E**) Pathways identified through significant upregulation of mRNA in KCB condition Day 3 versus Day 1. **(F**) Volcano plot of DEGs in D2 KCB condition compared to D2 KC condition. Log2 FC indicates the mean expression level for each gene. Each dot represents one gene. **(G)** Pathways identified from upregulated DEGs in D2 KCB versus D2 KC. **(H)** Differentially accessible peaks (DAPs) identified in D2 KCB versus D2 KC. **(I)** Pathways identified from DAPs as more accessible in KCB at D2. **(J**) Binding motifs identified as more accessible in D2 KCB versus D2 KC ADM. Statistical analysis performed using either a t-test of the area under the curve or one-way ANOVA. The padj for pathway analysis was calculated using enrichr for 2020_MSigDB_hallmark gene sets. \**p*<0.05 \*\**p*<0.01 \*\*\**p*<0.001 \*\*\*\**p*<0.0001 Scale bar represents 100μm.

During the ADM process, cells undergo large epigenetic and transcriptional changes as genes involved in maintaining an acinar cell identity are suppressed and genes that drive a ductal cell fate are induced. As PP2A is known to regulate the activity of chromatin modifiers and transcription factors (26), we wanted to determine if the accelerated transdifferentiation seen with the loss of B56α was due to large-scale changes in ADM associated gene programs. First, we performed RNA-seq on mRNA isolated from ADM structures in the KC and KCB^hm/hm^ conditions at 24H (1d), 48H (2d), and 72H (3d). In agreement with the qRT-PCR data and transdifferentiation from an acinar to a ductal cell identity, large changes in gene expression occurred in KC cells between 24H and 72H, with 1441 genes increased and 1626 genes decreased (p-adj<0.05, FC>1.5). In the KCB^hm/hm^ cells, even larger changes in gene expression occurred between 24H and 72H, with 2044 genes increased and 2545 genes decreased (**Fig 4D, S2E**). The differentially expressed genes at 72H generally showed good correlation between KC and KCB^hm/hm^ (**Fig S2F**), indicating that the overall transdifferentiation process was not dramatically altered; however, 944 genes were increased only in KCB at 72H (**Fig 4D**). This gene set was significantly enriched for MYC and E2F targets, consistent with the role of B56α in the negative regulation of MYC signaling (**Figure 4E**). In addition, there were differentially regulated genes (DEGs) between the genotypes at each time point, with the largest number of DEGs observed in KCB^hm/hm^ structures compared to KC structures at 48H (**Fig 4F**). These 302 genes were enriched for IFNγ response genes, pancreas beta cells, and KRAS signaling (**Fig 4G**). To identify sites of potential transcriptional activity underlying these gene expression changes, we performed ATAC-Seq on KC and KCB^hm/hm^ ADM structures at 24H, 48H, and 72H. We used MACS to define peaks for 2-3 biological replicates from each condition and observed good overlap of peaks (**Fig S2G**). We observed between 16,000-35,000 peaks for each sample, with the largest number of peaks and an overall increase in accessibility for both genotypes at 48H (**Fig S2H**). There were 1851 peaks with greater accessibility in KCB^hm/hm^ at 48h (**Fig 4H**). Taking the nearest associated gene, these areas of accessibility were associated with genes enriched in pathways such as epithelial to mesenchymal transition (EMT), inflammatory response, KRAS signaling up, and TGF signaling (**Figure 4I**). Furthermore, the enriched motifs identified using HOMER were associated with developmental transcription factors (Prdm1, Hoxc13, Nkx2-3), hematopoietic transcription factors (Tcf1, Klf1) downstream regulators of TGF-beta signaling (Smad3, Smad5), and inflammation (Stat3, Irf4, Irf3, Irf7) (**Fig 4J**). Together, the RNA- and ATAC-seq indicate that the loss of B56α intensifies KRAS and MYC signaling, and potentially drives a more dedifferentiated cell state that increases the potential of acinar cells to undergo ADM in response to activated oncogenic KRAS signaling.

### Loss of PP2A-B56α accelerates KRAS^G12D^ PDAC initiation *in vivo*

The pancreas-specific expression of KRAS^G12D^ faithfully recapitulates pancreatic cancer initiation, including ADM and the formation of PanINs *in vivo* (4,5). This progression is marked by a loss of acinar tissue, formation of mucin producing PanIN lesions, and stromal expansion. To determine if the loss of B56α can promote PDAC initiation *in vivo*, pancreata from WT, B56α^hm/hm^, KC, and KCB^hm/hm^ were harvested at 2-month and 5-month timepoints and evaluated for changes associated with early PDAC progression. B56α^hm/hm^ mice display normal pancreatic histology, indicating that the loss of B56α alone is not sufficient to drive PDAC initiation. However, similar to our *ex vivo* ADM findings (**Fig 4**), there was a reduction in healthy acinar area starting at 2 months of age in KCB^hm/hm^ mice compared to KC mice, with a significant loss by 5 months (**Fig 5A-B**). We then stained pancreatic tissues with alcian blue, a basic dye that positively labels acidic patches of mucin expression within PanIN lesions, to determine the impact on progression. Loss of B56α resulted in accelerated PanIN formation, with KCB^hm/hm^ mice displaying a significant increase in alcian blue positive lesions at both 2 and 5 months of age compared to KC tissues (**Fig 5A, C**). These aberrant morphological changes within KCB^hm/hm^ tissues were accompanied by increased collagen deposition, a hallmark of PDAC progression (27), as indicated by masson’s trichrome (MTC) (**Fig 5A, D**). Together, the decrease in healthy acinar area, increase in PanIN lesions, and increase in collagen deposition suggest that loss of B56α cooperates with KRAS^G12D^ to accelerate PDAC initiation and the development of pre-malignant lesions.

**Figure 5:**
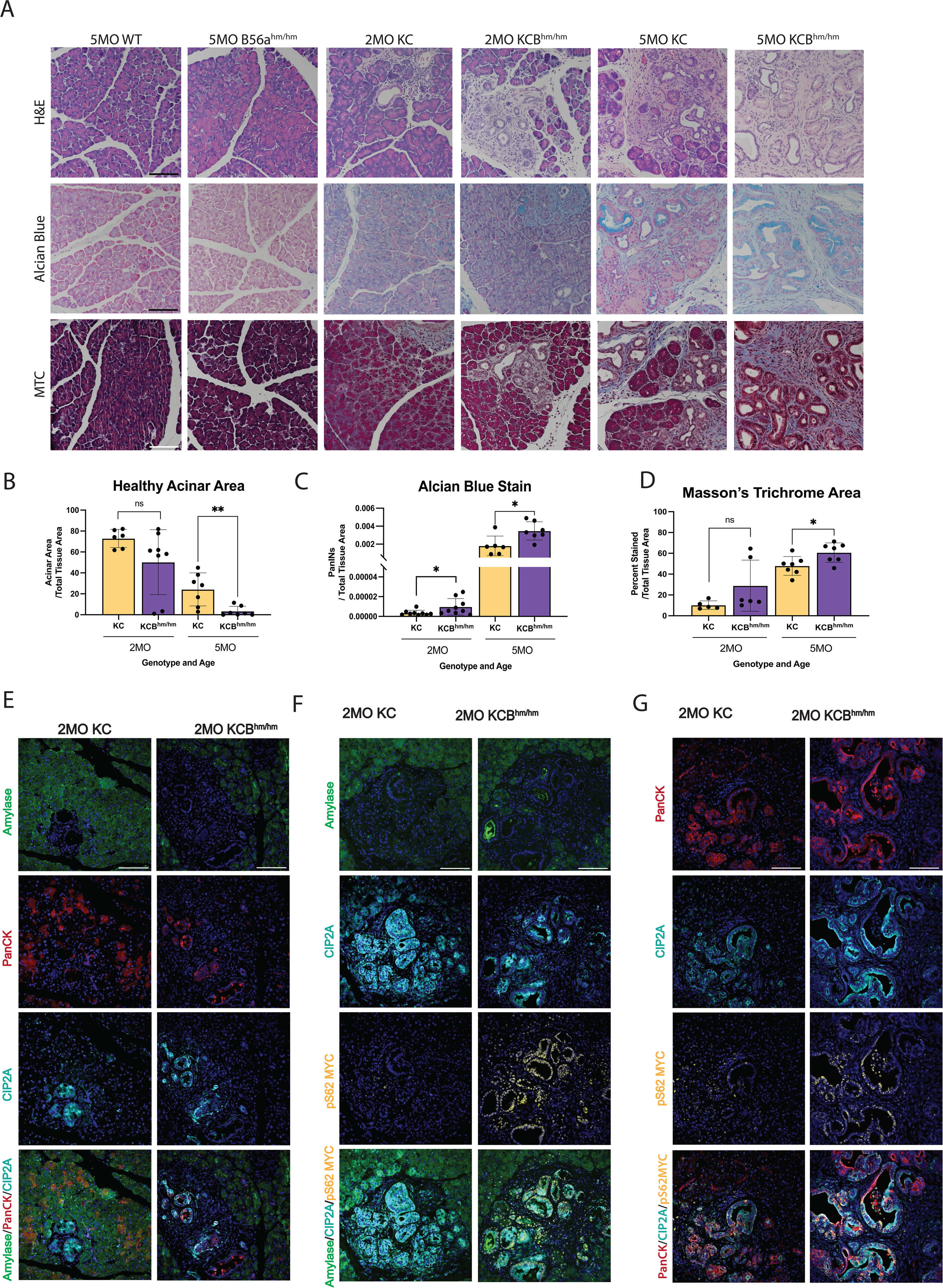
Loss of B56a promotes progression of PDAC precursor lesions. **(A)** Representative 2MO or 5MO timepoints from pancreatic tissue in respective genetic mouse models stained with H&E (top), Alcian Blue (middle), and Masson’s Trichrome (bottom). **(B-D)** Quantifications of healthy acinar area, stromal content, and number of PanINs quantified from stains represented in A, respectively, at 2MO and 5MO timepoints. All analysis represented at least 6 biological replicates per condition. **(E)** Amylase, CIP2A, pS62 MYC, and merged immunofluorescent stains in pancreata harvested 2MO KC or KCB^hm/hm^ mice. **(F)** PanCK, CIP2A, pS62 MYC and immunofluorescent stains merged in pancreata harvested 2MO KC or KCB^hm/hm^ mice. All images represent at least three independent biological replicates. \**p*<0.05 \*\**p*<0.01 n.s. = not significant Scale bar represents 100μm.

Based on our findings that KRAS^G12D^ increases the expression of CIP2A and reduces B56α activity (**Figure 1**, **2**), we analyzed CIP2A and MYC expression within acinar (Amylase+) and ductal (Pan-Cytokeratin (PanCK+)) populations (**Fig 5E-G, Fig S3A-C**). CIP2A expression was strongly correlated with PanCK+ cells in both KC and KCB^hm/hm^, highlighting a role for PP2A-suppression in promoting the ductal identity. Low level expression of pS62 MYC was observed in both acinar and ductal populations of 2-month KC tissues, which increased in PanIN lesions by 5 months (**Fig 5F-G, S3B-C**). In contrast, the loss of B56α led to high pS62 MYC expression in both acinar and ductal populations at earlier timepoints, with 2-month KCB^hm/hm^ tissues displaying higher pS62 MYC levels than KC tissues at 5-months of age. Overall, these results suggest that PP2A-B56α plays an important role in restraining pancreatic cellular plasticity during PDAC initiation and the suppression of this subunit significantly contributes to oncogenic KRAS PDAC progression.

### Therapeutic Reactivation of PP2A attenuates acinar-to-ductal transdifferentiation

Since the active PP2A holoenzyme requires the association of three individual subunits (A, B, and C), genetic strategies to model B56α activation are extremely limited. Therefore, we took advantage of the small molecule activator of PP2A (SMAP), DT061, to determine if PP2A activation can prevent acinar cells from undergoing ADM. This compound binds at the interface of all three PP2A subunits (A, B, and C) and functions to nucleate the PP2A holoenzyme (28). This small molecule PP2A modulator has been shown to preferentially promote the stabilization of PP2A-B56α heterotrimers. Acinar cells isolated from KC pancreata were plated in an ADM assay and treated with either DMSO control or increasing doses of DT061 for 72H (**Fig 6A**). Endpoint cultures were then quantified for acinar or ductal-like structures. Our data show that DT061 reduced the percent of duct-like structures in a dose-dependent fashion (**Fig 6B**), and the ductal structures that did form were significantly smaller compared to control (**Fig 6C, D**). To validate that the phenotypic loss of ductal-like cells correlates with a loss of ductal identity, we isolated RNA from the endpoint structures in each condition and ran qPCR for ductal identity markers. We found that increasing doses of DT061 led to lower levels of canonical ductal markers, *Ck19* (**Fig 6G**). DT061 also decreased mRNA levels of *Myc* and *Cip2a* (**Fig 6E, F**) and increased B56α (*Ppp2r5a*) (**Fig 6H**). Together these findings suggest that PP2A activation antagonizes ductal differentiation.

**Figure 6:**
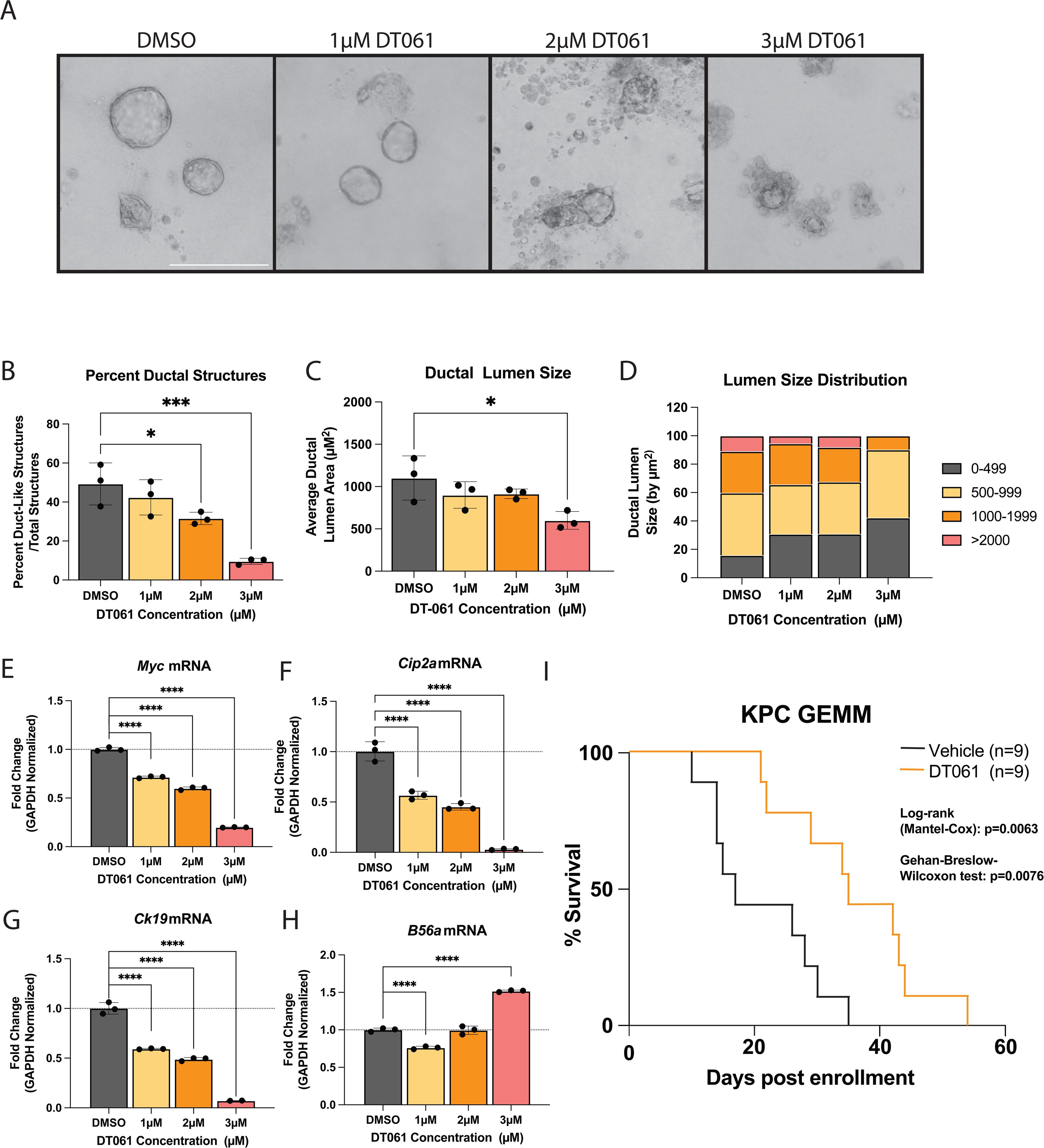
Re-activation of PP2A-B56a prevents initiation of pancreatic cancer. **(A)** Representative images of acinar to ductal differentiation in KC acinar cells after 72H in collagen and treated with increasing doses of SMAP, DT061. **(B)** Percent duct-like structures over total structures present at each concentration of DT061 treatment. **(C)** Average ductal lumen size of duct-like structures in each condition. **(D)** Distribution of the ductal lumen measurements at each concentration of DT061. **(E-H)** qPCR of mRNA isolated from the organoids in collagen after 72H of DT061 treatment at each concentration dosed. ADM data representative of one biological replicate. Assay was repeated with same trends in three independent biological replicates. **(J)** Survival curve of *PDX1-Cre; P53^+/-^; LSL-KRAS^G12D^* (KPC) mice enrolled in a survival experiment when 100mm^3^ tumor was identified and then treated twice daily with 15mg/kg DT061 via oral gavage or with vehicle until mice succumb to PDAC disease burden. \**p*<0.05 \*\*\**p*<0.001 \*\*\*\**p*<0.0001 Scale bar represents 100μm.

Recent studies have demonstrated that inhibition of mutant KRAS with the small molecule, MRTX1133, results in strong tumor suppression in KPC allografts, highlighting the importance of this pathway to PDAC survival and progression (29). As KRAS^G12D^ inhibited PP2A function and increased oncogenic MYC levels, (**Fig 1**) and activation of PP2A with DT061 attenuated KRAS^G12D^-driven phenotypes, we next sought to investigate whether DT061 could suppress late-stage PDAC tumor growth. For these studies, we utilized the KPC (*LSL-KRAS^G12D/+^; P53^-/+^; PDX1-Cre)* PDAC mouse model (30). These mice represent a highly aggressive genetic mouse model of PDAC that accurately reproduces several aspects of human disease, including desmoplasia and metastatic disease, and is therapeutically refractory to most standard-of-care chemotherapies(31). We monitored KPC mice weekly by high-frequency ultrasound, and then randomly enrolled mice into treatment arms (Vehicle vs DT061) when tumors reached 5-7mm in diameter. DT061 significantly increased the overall survival of KPC mice compared to vehicle treated animals (**Fig 6I**). Together, these results suggest that therapeutic re-activation of PP2A can antagonize both the initiation and the progression of PDAC tumors *in vivo* in the context of key oncogenic drivers such as KRAS.

## DISCUSSION

In this study, we have determined a mechanism by which oncogenic KRAS signaling inhibits PP2A-B56α activity to promote both the initiation and progression of pancreatic cancer. Our data demonstrate that activated KRAS signaling promotes the upregulation of the PP2A-B56-specific endogenous inhibitor, CIP2A, which attenuates B56α binding to the PP2A complex. Further, we demonstrate that PP2A-B56α negatively regulates the progression of KRAS^G12D^-driven ADM and the initiation of pancreatic precursor lesions *in vivo*. Previous studies have demonstrated that transcriptional deregulation of MYC can accelerate KRAS^G12D^ PDAC tumor development and is necessary to maintain tumor growth (32). However, MYC is known to be tightly regulated at the transcriptional, translational, and posttranslational levels. Here, we demonstrate that the inhibition of B56α increased the expression of the active form of MYC (pS62) during PDAC progression, implicating the posttranslational regulation of MYC as critical mechanism of PDAC tumor suppression. These findings also indicate that the contribution of aberrant MYC signaling in the absence of transcriptional alterations may be grossly underestimated in PDAC patients.

A wide range of mechanisms have been implicated to contribute to increased MYC signaling downstream of KRAS, including those that increase the transcription and/or stabilization of CIP2A. Studies have shown MYC, E2F1, and ETS1 can drive CIP2A transcription downstream of RAS, and KRAS-induced expression of SURVIN can protect CIP2A from autophagy/lysosomal dependent degradation (33–35). Further, CIP2A is stabilized through its interaction with PP2A (21). Our studies support the notion that CIP2A transcription and translation are regulated through the activation of KRAS, as KRAS^G12D^ induction led to a significant upregulation of CIP2A expression. Further, we show that KRAS^G12D^ results in the disassociation of B56α with the PP2A catalytic subunit in a CIP2A dependent manner and leads to a simultaneous increase in MYC protein levels. Using crosslinking combined with mass spectrometry with purified proteins, Pavic et al. have recently proposed a new mechanism by which CIP2A replaces the PP2A-A subunit from the PP2A heterocomplex and prevents B56-subunits from binding their target (36). While these studies highlight novel mechanistic regulation that occurs between CIP2A and PP2A, further studies are needed to determine the influence of CIP2A on B56 disassociation rates from the PP2A-A and C subunits and the potential for differential regulatory mechanisms of individual B subunits and in cancer versus normal cells.

Using *ex vivo* implantation of acinar clusters to measure ADM, we demonstrated that the acceleration of transdifferentiation by loss of B56α is a cell intrinsic process. At endpoint, KCB^hm/hm^ cells that have undergone ADM display an enrichment of MYC, E2F, and cell cycle gene programs. We found similar results *in vivo*, with KCB^hm/hm^ mice displaying increased pS62 MYC in early ADM lesions. These results indicate that B56α plays a critical role in the posttranslational regulation of MYC during early PDAC development. The loss of B56α also led to a specific enrichment of transcriptional and epigenetic gene programs associated with increased KRAS signaling. These findings are consistent with PP2A negatively regulating downstream KRAS effectors and the recently published data that RAS and PP2A signaling pathways converge at co-regulation of epigenetic modifiers (37). In addition to KRAS and MYC, we found chromatin reorganization near genes implicated in ADM and/or PDAC development, including EMT, TGF-β signaling, and inflammation, as well as an enrichment of transcription factor (TF) binding motifs associated with cell plasticity. Studies have shown that knockdown of the PP2A-A subunit enriches for these same pathways (37), indicating that the B56α subunit is the primary mechanism through which this regulation occurs and further supporting a role for PP2A in maintaining normal cellular homeostasis. PP2A has also been shown to suppress cell-intrinsic inflammatory signaling (38–40). Therefore, loss of B56α may contribute to and further exacerbate the inflammation that drives the process of ADM and PDAC initiation. Overall, our understanding of the posttranslational regulation of epigenetics by PP2A is still in its infancy. Therefore, further insight into how B56α alters epigenetic regulation of ADM is of critical importance.

PP2A is a unique tumor suppressor as it is rarely genetically deleted or mutated in most human tumor types. Instead, loss of PP2A activity is primarily driven through suppression by endogenous inhibitors like CIP2A (9). Since the ability of PP2A to form a functional holoenzyme remains intact, the therapeutic re-activation of this phosphatase has shown strong preclinical responses in several cancers, including prostate, ovarian, breast and lung cancers (41–45). Furthermore, activation of PP2A, either through genetic or pharmacological means, functions synergistically with various kinase inhibitors implicating PP2A as a critical rheostat of therapeutic response (16,37,46). Our studies show that DT061 not only abrogates the process of ADM, but also improves long-term survival of spontaneously driven PDAC in KPC mice when treated with small molecule modulators of PP2A. Since DT061 preferentially promotes the activation of B56α containing PP2A heterotrimers, our results strongly support a critical role for this subunit in *in vivo* tumor progression and support further investigation into the mechanisms by which therapeutic re-activation of PP2A regulates late-stage disease (28). Additionally, KRAS inhibitors have shown excellent pre-clinical efficacy in PDAC, but re-activation of KRAS signaling has emerged as a significant resistance mechanism in multiple tumor types (47–50). Our studies implicate PP2A as a critical negative regulator of KRAS signaling and, therefore, the combination of KRAS inhibitors and PP2A activators may represent a novel therapeutic strategy to attenuate acquired resistance.

Finally, our studies represent one of the few *in vivo* experimental models identifying a role for a specific PP2A B subunit in regulating tumor development and progression. Redundancy between PP2A subunits has been a confounding factor in the field and potentially masks the contribution of specific PP2A complexes, like PP2A-B56α, to tumor function. It is unknown if loss of one B subunit drives the incorporation of another into the PP2A complex. Therefore, it is essential that future studies analyze the dynamic interactions between B subunits within tumors. However, given that each PP2A B subunit potentially has unique and context dependent functions, the development of therapeutic compounds that leverage individual subunits would be an extremely impactful clinical advance.

## METHODS

### Cell Culture

iKRAS #3(51) cells were gifted from Dr. Marina Pasca di Maglinao (University of Michigan, Ann Arbor, MI). iKRAS #3 cells were cultured in RPMI1640 + 10% fetal bovine serum. iKRAS #3 cells were maintained in 1ug/mL doxycycline (dox). HPDE-iKRAS cells were gifted from Dr. Gordon Mills (OHSU, Portland, OR) and cultured in Keratinocyte Serum-Free Media (KSFM) supplemented with epidermal growth factor and bovine pituitary extract (Fisher Scientific, 17-005-042). Virus to construct stable HA-B56α overexpression cell lines were produced using HEK293T cells transfected using Lipofectamine 3000 (Fisher Scientific, L3000015) with the plasmid of interest (POI) and two plasmids encoding packaging proteins for viral vector (pAX.2 Addgene Plasmid #35002), pMD2.G Addgene Plasmid #12259). Parental cells were pre-treated with 10μg/mL polybrene for 1 hour before adding viral particles to media. Media was changed after 24H and selected using antibiotic matching POI. Cells transfected with siCIP2A (Horizon Discovery, L-014135-01-0005) were transiently transfected using Lipofectamine 3000 and then utilized 72H after transfection. All cell lines were routinely tested for *Mycoplasma* using PCR-based strategies and grown at 37°C in a 5% CO2 atmosphere.

### Quantitative RT-PCR

RNA was isolated using the ThermoScientific GeneJET Purification Kit (Thermofisher, K0732). Eluted RNA concentration was measured using the BioTek Synergy H1. Reverse transcription PCR was performed using the High-Capacity cDNA Reverse Transcription Kit (Fisher Scientific, 43-688-14). Quantitative PCR is performed on the QuantStudio 3 using SYBR Green (Fisher Scientific, A25743) and the primers listed on the table below made custom from Integrated DNA Technologies. The relative fold change relative to vehicle was analyzed using the ΔΔ(*C*t) method.

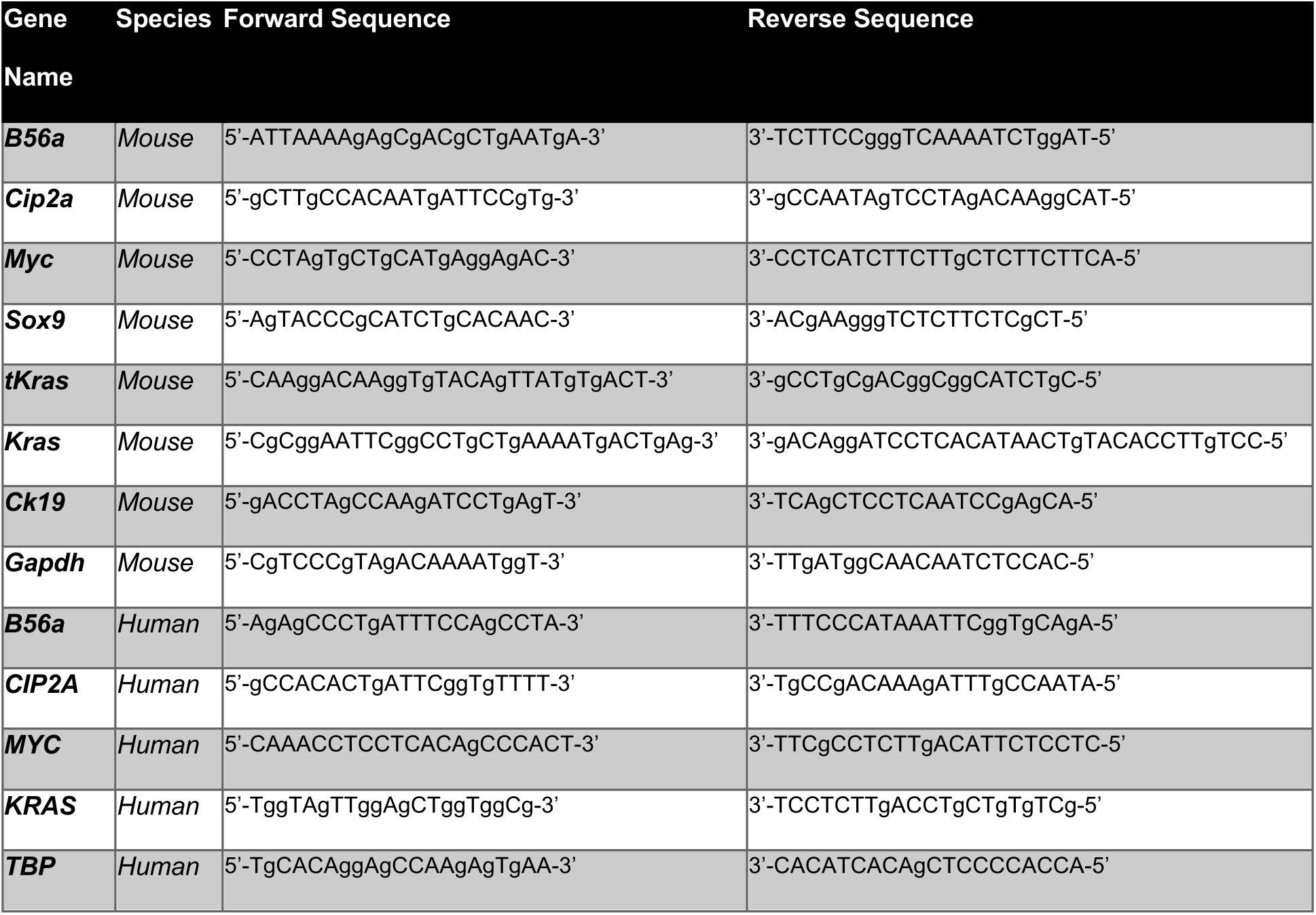

### Western Blot Analysis

Cells were lysed (20mM Tris pH7.5, 50mM NaCl, 0.5% Triton X-100, 0.5% deoxycholate, 0.5% SDS, and 1mM EDTA) and protein concentration was measured using the DC protein assay kit (Bio-Rad,5000112) and absorbance was read on the Synergy H1. Lysate was then treated with SDS sample buffer (57.5% glycerol, 125mM Tris pH 6.8, 5% SDS, 5% b-mercaptoethanol, bromophenol blue) and boiled at 95°C for 5 minutes. SDS page was run on 4-15% gradient protein gels using the Criterion Electrophoresis Cell (Bio-Rad, 1656001) and transferred to a PVDF membrane using the Trans-Blot semi-dry transfer system (Bio-Rad, 1704150). Membrane was blocked in Licor TBS blocking buffer (Fisher Scientific, NC1660550) for 1H at RT and then incubated with primary antibody overnight at 4°C. Membranes were then incubated with secondary antibody for 1H at RT and scanned using the Licor Odyssey DLx Imaging system and analyzed using Image Studio software.

| Gene Name | Catalog Number | LOT Number | Company |
| --- | --- | --- | --- |
| KIAA1524 | HPA039570 | 000008181 | Atlas Biosciences |
| CIP2A | sc-80659 | H2522 | Santa Cruz Biotechnologies |
| B56a | C Terminal Clone 3A6D3 | N/A | Egon Ogris Lab |
| MYC | ab32072 | GR3377350-8-YR091119PS | Abcam |
| GAPDH | M4300 | 820845 | ThermoFisher |
| Phospho-ERK 1/2 (pS42, pS44) | 9101 | 31 | Cell Signaling Technologies |
| ERK 1/2 | 4695 | 28 | Cell Signaling Technologies |
| IRDye 680RD Goat anti-Rabbit | 926-68071 | D20809-05 | Licor |
| IRDye 800CW Goat anti-Mouse | 926-32210 | D20802-25 | Licor |

### Proximity Ligase Assay (PLA)

The PLA was performed using the Duolink In-Situ Red Starter Kit (Millipore Sigma, DUO92101) following manufacturers instructions with an extended amplification period of 3H instead of the recommended 1H. Treated cells were then counterstained with DAPI for 5 minutes and ActinGreen 488 (Thermofisher, R37110) for 20 minutes at RT. Cells were imaged using the Nikon Eclipse Ni upright fluorescent microscope. Fluorescent images were gated in FIJI and PLA puncta per cell were quantified and analyzed using a one-way ANOVA.

### Soft Agar Assay

To perform this assay, a 12 well plate was coated 500uL with a 1:1 mixture of 1.4% noble agar solution mixed and 2X RPMI media supplemented with 20% FBS. Then 40,000 cells/well were resuspended in 2X RPMI+20% FBS media and mixed 1:1 with a 0.7% noble agar solution. 500uL of the cell suspension was plated per well and allowed to solidify at RT. Finally, the cells were topped with 1mL 1XRPMI +10% FBS per well and treated with a final concentration of 1ug/mL dox. Cells were allowed to grow for two weeks and dox was re-dosed every 3-4 days. Colonies were measured by percent area covered using FIJI and conditions were statistically analyzed using a student’s t-test between conditions.

### Clonogenic Assay

One thousand cells/well were seeded in a 6-well plate. HPDE-iKRAS cells and iKRAS #3 cells were supplemented with doxycycline every 48-72H (50ng/mL and 1μg/mL, respectively). Cells were allowed to grow for six to seven days and then fixed and stained with 0.1% crystal violet solution (0.1% crystal violet, 20% methanol, 80% water) for 2H at RT. Then, crystal violet-stained cells were rinsed with PBS and water, plate was allowed to air-dry overnight. For imaging, whole-well images of the plate were scanned on the EVOS M7000 and percent area covered by crystal violet-stained cells was quantified using FIJI and analyzed using student’s t-test.

### Mouse Strains

All animal studies were performed in compliance with Purdue University (West Lafayette, IN) animal use guidelines after approval by the Purdue Institutional Animal Care and Use Committee. *NOD.Cg-Rag1^tm1Mom^ Il2rg^tm1Wjl^/SzJ* (NRG)(Jackson Laboratory #007799) mice were used for allograft studies. PDAC initiation and progression were analyzed using *PDX1-Cre; LSL-Kras^G12D^* (KC) and *PDX1-Cre; LSL-Kras^G12D^; B56^hm/hm^* (KCB^hm/hm^), *B56^hm/hm^, and WT* genetically modified mice on a 129S1/SvlmJ and C57/BL6 strain mixed background (129J/Bl6). For timepoints, both male and female mice were euthanized at 2M and 5M months of age by CO2 followed by cervical dislocation. Pancreatic tissue was then formalin fixed and paraffin embedded. Mouse genotypes were confirmed by clipping ears and utilizing the HOTSHOT DNA isolation protocol (Truett et al. *BioTechniques*, 2000) to extract DNA. PCR primers in the table below were used for gene identification.

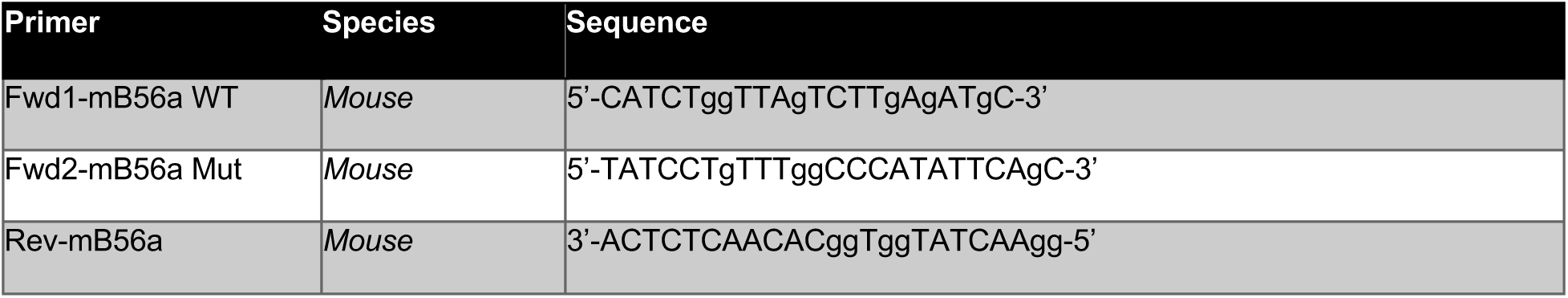
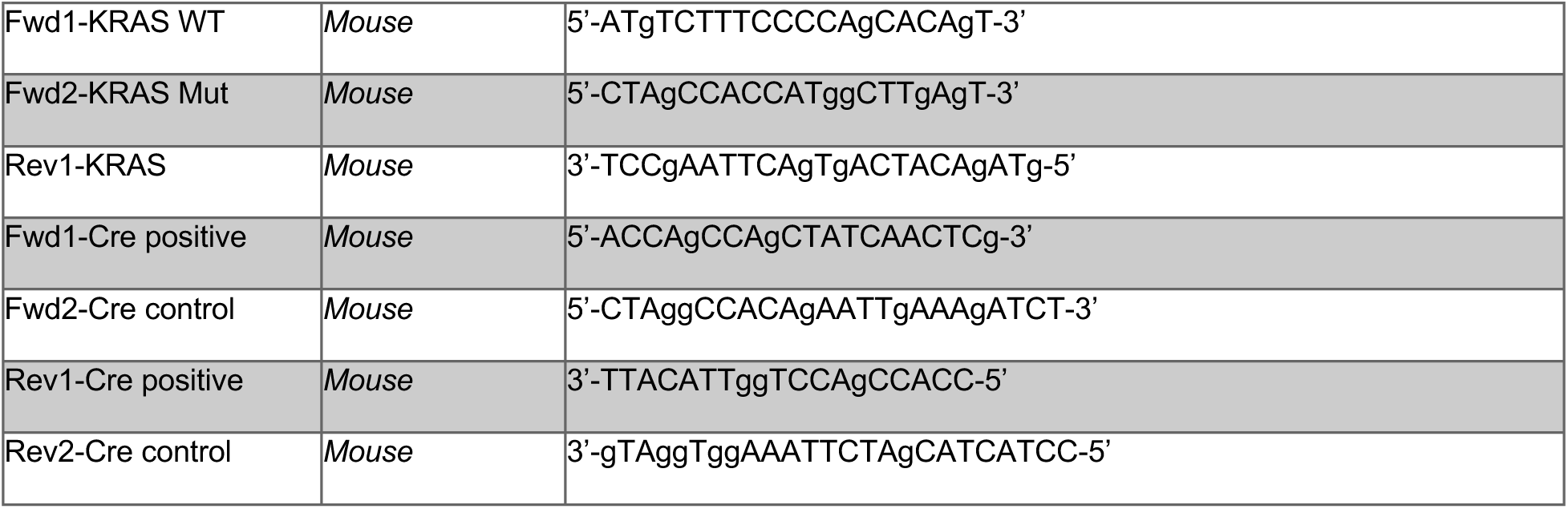

### Allograft Studies

Orthoptopic injections were performed as previously described(52). Briefly, mice were treated with 2mg/mL meloxicam and anesthetized with 2-3% isoflurane supplemented with 1g/L oxygen. An incision was made just below the ribcage on the left side. Spleen was removed from abdomen with the pancreas following. iKRAS #3 stable cell lines (EV, B56αOE) were mixed 1:1 with serum-free RPMI and Matrigel and 5,000 cells were injected into the pancreas. Pancreas was placed back into the abdomen, the muscle layer was sutured, and skin was closed using staples. Mice were monitored for recovery to sternal recumbency and monitored daily post-surgery until staples were removed after one week. Tumors were then measured weekly using the VEVO3100 rodent ultrasound machine and measured using Vevolab software.

### Acinar-to-ductal metaplasia assays

ADM Assay protocol was adapted from Martinez and Storz. *JoVE,* (2019). The collagen matrix was prepared consisting of 3.5mg/mL type I rat tail collagen, 1X Waymouth Media, and 7.7mM NaOH. 500µL of collagen matrix was added to the bottom of each well in a 12 well plate. Primary mouse acinar cells were obtained from KCB^hm/hm^, KC, B56^hm/hm^, and WT mice of 129J/Bl6 background. Mice were euthanized at 4-6 weeks by isoflurane induction followed by cervical dislocation and subsequent cardiac puncture to endure death. 2-3 Pancreata per genotype were collected on ice in HBSS + 1% Pen/Strep and 100ng/mL Soybean Trypsin Inhibitor (STI). Pancreata were processed into small pieces using scalpels and then placed into 2mg/mL Collagenase I supplemented with 100ng/mL STI. Pancreata were incubated in collagenase solution for 20-30mins shaking at 37°C until small acinar clusters of approximately 100µm or less were visible under the microscope. Pancreatic tissue was broken up by gentle 1mL pipetting. Acinar cells were then pelleted at 900xg for 2mins at 4°C, removed from collagenase and washed with a 5%FBS/1%PenStrep/HBSS solution three times. Cells were then resuspended and passed through a 100µM filter to remove debris. Acinar cells were layered slowly on top of a 30%FBS/HBSS solution and separated by slow centrifugation at 200xg for 2mins at 4°C. FBS/HBSS solution was removed, and cells were resuspended and diluted in ADM Assay Media (1X Waymouth Media, 1% FBS, 100ng/mL STI, 1ng/mL Dexamethasone, and 1% Pen/Strep) until approximately 20 structures/10X field were observed in a 96 well plate. Acinar clusters in ADM assay media were mixed 1:1 with collagen matrix and 500µL of solution plated per well in a 12 well plate. Plate incubated at 37°C until collagen matrix solidified, and wells were topped with 500µL of ADM Assay Media. For ADM Assays testing DT061 efficacy in ADM, only KC genetic mice were used, and media was supplemented with DMSO or increasing doses of DT061. The acinar to ductal transdifferentiation process was observed using an EVOS M7000 through timelapse imaging every 6 hours for 72 hours total in an on-stage incubator maintaining 5% CO2 and 37°C. For all replicates, 400 structures were measured per timepoint (1d, 2d, 3d post-implantation). For the kinetics assay, 100 acinar structures per well, 3-4 wells/condition were followed throughout the 72H timelapse images and marked when transdifferentiated into duct-like structures using FIJI.

### RNA-Seq processing

RNA was isolated using the ThermoScientific GeneJET Purification Kit (Thermofisher, K0732). Eluted RNA concentration was measured using the Synergy H1. mRNA was polyA enriched and RNA-seq libraries were generated and Illumina sequenced by Novogene.

### RNA-Seq analysis

Datasets were trimmed using fastp (v0.23.2) (53) to remove adapter sequences and bases with a Phred quality score lower than Q30. Remaining reads with lengths greater than 50bp post-trimming were retained for further analysis. Trimmed reads were aligned to the Mus musculus reference genome (GRCm39) from Ensembl release 109 using STAR Aligner (54) version 2.7.4a in two-pass mode. Reads were assigned to features using featureCounts (55) version 1.6.1 in paired-end mode and using the ‘unstranded’ quantification approach. The feature count matrix was imported into Partek Flow software (v10.0.23.0720). The counts were filtered for low-expression genes and DEGs were generated using DESeq2 (56) (Partek Flow v10.0.23.0720) default parameters. An adjusted pvalue<0.05 and fold change>1.5 was used for downstream DEG analysis. PCA plots were generated using normalized count files.

### ATAC-seq

KC and KCB^hm/hm^ cells were cultured in ADM assays as above. Cells were harvested at 24, 48 and 72h after plating through digestion by 2mg/mL Collagenase I for 20-30 minutes. Isolated cell clusters were then centrifuged, washed twice with 1XPBS, and frozen in 5% DMSO as single cell suspensions. Cells were thawed and 50,000 cells were counted and processed according to the Omni-ATAC protocol (Corces et al. *Nature Methods* 2017). The final libraries were subjected to double size selection according to a published protocol (Harmacek et al. *Methods Mol Bio* (2018) and quality checked using Qubit and Agilent TapeStation by Purdue Genomics Core Facility. Equimolar libraries were pooled using cluster numbers generated using Miseq (Purdue Genomics Core Facility). Libraries were sequenced using 150bp PE sequencing on NovaSeq 6000 platform (Novogene, Sacramento, CA). All experiments were performed in three biological replicates.

### ATAC-seq analysis

Data processing and analysis were done using Partek Flow software (v10.0.23.0531). Raw reads were trimmed for Nextera adapter (CTGTCTCTTATACACATCT), minimum length>20 and quality>20 using Cutadapt_4_2 (57). Trimmed reads were aligned to mouse genome mm10 build using BWA-MEM (v0.7.17) (58) with default parameters. Alignments were filtered for mitochondrial reads, PCR duplicates, MAPQ<20, singletons, not primary reads, blacklisted regions and chrY alignments to keep properly paired alignments for downstream steps. The filtered alignments were used for peak calling using MACS (3.0.0a7) (59) in 2 ways: ATACseq-mode wherein peaks were called in samples individually, and ChIPseq-mode wherein peaks were called by comparison between conditions (biological replicates were pooled for comparison). Volcano plot shows peaks with adjusted pvalue<0.05 and fold change>2. Filtered alignments were used for BigWig generation using deeptools (60) bamCoverage with --normalizeUsing CPM option. Homer was used for motif analysis with -size 200-mask options. PCA plots were generated using CPM normalized filtered alignments.

### Bioinformatics and Data Visualization

ATAC-Seq Peak overlaps were performed using Bedtools Intersect Intervals (v2.30.0) (61)in the Galaxy interface (Nucleic Acids Research, Volume 50, Issue W1, 5 July 2022, Pages W345– W351, doi:10.1093/nar/gkac247). Peaks were annotated and assigned to the nearest gene using the mm10 genome and the ChIPseeker (62) annotate software package in R. Gene overlaps were calculated using the InteractiVenn online interface (63). Gene and peak overlaps were visualized using eulerr software package in R. Heatmaps and metagene analyses of ATAC data were performed in the Galaxy interface using DeepTools (v3.5.2) computeMatrix and plotHeatmap. Pathway analysis was performed using Enrichr (Maayan Lab cloud) (64) for the MSigDB Hallmark 2020 gene sets. Volcano plots and pathway enrichment were plotted using Prism 10.

### Tissue Immunofluorescence

Slides incubated twice in xylene for 10 minutes and then rehydrated by incubation in 100% EtOH I, 100% EtOH II, 95% EtOH, 70% EtOH, and 100% PBS for 5 minutes each. Slides were then cooked in a pressure cooker at high pressure for 10 minutes submerged in 1X High pH Antigen Retrieval Solution first (10 mM Tris, 1 mM EDTA solution, pH 9.0), cooled to RT for 30min, rinsed in PBS, then cooked in the pressure cooker for an additional for 10mins in 1X Low pH Antigen Retrieval (10mM Sodium Citrate, pH 6.0). Slides were cooled to room temperature for 30 minutes, then rinsed with PBS 3 times for 3 minutes each. Slides were blocked in 3% BSA at RT for 1 hour. Primary antibodies were diluted in 3% BSA with mouse-on-mouse kit (Vector Laboratories, MKB-2213-1), and incubated at 4°C overnight. PBS-Tween (0.025%) was used to wash slides 3 times for 5 minutes each. Secondary antibody diluted to 1:200 in 3% BSA, incubated in the dark for 1 hour at room temperature. After secondary incubation, the slides were washed with PBS-Tween (0.025%) 3 times for 5 minutes each. Slides were incubated with DAPI at 1:1000 (Sigma-Aldrich, D9542) for 5 minutes, the excess solution was blotted off and coverslip was mounted using Prolong Gold Antifade Mountant (Invitrogen, P36934) and tissue was imaged on Nikon Eclipse Ni upright microscope and analyzed using FIJI.

| Gene Name | Catalog Number | LOT Number | Company |
| --- | --- | --- | --- |
| **KIAA1524** | HPA039570 | 000008181 | Atlas Biosciences |
| **pS62 MYC** | 4B12 | N/A | Rosalie Sears Lab (65) |
| **Amylase** | A8273-1VL | 0000121533 | MilliporeSigma |
| **Anti-pan Cytokeratin** | sc-8108 | J0721 | Santa Cruz Biotechnologies |
| **Chicken anti-Rat IgG**<br>**(H+L), Alexa Fluor 488** | A21470 | 2331905 | Invitrogen |
| **Goat anti-Mouse IgG**<br>**(H+L), Alexa Fluor 647** | A21235 | UK290265 | Invitrogen |

### Immunocytochemistry

Cells were fixed using 4% paraformaldehyde for 15 minutes at RT, washed with PBS 3 times for 5 minutes, then permeabilized using 0.3% triton-x100 for 25 minutes at RT. Cells were then stained using primary antibody diluted in 3% BSA and incubate overnight at 4°C. Cells were washed in PBST (0.025%) 3 times for 5 minutes and then stained with secondary antibody diluted in 3% BSA in the dark for one hour at RT. Cells were washed 3 times in PBST for 5 minutes. Cells were incubated with DAPI at 1:1000 (Sigma-Aldrich, D9542) for 5 minutes and Prolong Gold Antifade Mountant (Invitrogen #P36934) was added before coverslips were applied. Cells were imaged on Nikon Eclipse Ni upright microscope and analyzed using FIJI.

| Gene Name | Catalog Number | LOT Number | Company |
| --- | --- | --- | --- |
| KIAA1524 | HPA039570 | 000008181 | Atlas Biosciences |
| Goat anti-Rabbit IgG<br>(H+L), Alexa Fluor 594 | A32740 | WK333741 | Invitrogen |

### Alcian Blue Staining

The Alcian Blue (pH 2.5) Stain Kit (Vector Laboratories #H-3501) was used for this protocol. Slides were hydrated to distilled water; incubated in 100% xylene twice for 10 minutes, 100% EtOH twice for 5 minutes, 95% EtOH for 5 minutes, 70% EtOH for 5 minutes, and ddH2O twice for 5 minutes. Acetic Acid Solution was applied onto the tissue sections for 3 minutes, then the excess was tipped off. The Alcian Blue Solution was then applied to the tissue sections for 1 hour at 37°C. The slides were rinsed for 10-30 seconds in the Acetic Acid solution to remove the excess Alcian Blue Solution. The slides were rinsed for 2 minutes in running tap water followed by 2 changes of distilled water for 2 minutes each. Nuclear Fast Red solution was applied for 5 minutes. The slides were again rinsed for 2 minutes in running tap water followed by 2 changes of distilled water for 2 minutes each. Slides were dehydrated through graded ethanol and then mounted. Images were taken at 10x magnification and tiled together using an EVOSM7000 to calculate the number of PanINs in the total pancreas tissue area. The number of alcian blue positive panINs were counted using FIJI and normalized by dividing the total area of the pancreatic tissue section stained.

### IHC Tissue Staining

Tissue sections (4 μm) were hydrated as above and then stained with Hematoxylin 7211 (Thermo Fisher #7211) for 2 minutes then incubated in an ammonia-based bluing solution for 1 min. Slides were rinsed and then dehydrated to 95% ethanol. Then Slides were incubated in an Eosin Y counter stain (Fisher, 22-500-063) for 15 seconds. Slides were then dehydrated up to Xylene and allowed to dry at RT until Xylene was evaporated. Slides were cover slipped with Permount (Fisher, SP15-100) solution. For the Masson’s Trichrome Stain, the protocol outlined in Carson et al. *Chicago ASCP* (2015).

## ACKNOWLEDGEMENTS

We would like to thank the Pasca di Magliano lab (University of Michigan) for providing cell lines and historical histology slides for use; the Sears lab (OHSU) for providing cell lines, antibodies, mouse models, and technical support; and finally, the Narla lab (University of Michigan) for providing antibodies, DT061 stocks, and technical support for experiments. We would like to thank the Purdue Institute for Cancer Research (NIH grant P30 CA023168), the Purdue Histology Core, the Collaborative Core for Cancer Bioinformatics (C^3^B), and the Purdue Genomics Facility for their contributions to the data produced in this publication. We would also like to thank all the members of the BAP lab for editing of the manuscript and other helpful suggestions. G.N. acknowledges the support of the Rogel Cancer Center and is a Rogel Scholar. R.C Sears was supported by NCI U01 CA224012, R01 CA186241, and DoD PA210068 and the Brenden-Colson Center Foundation. S. L. Tinsley was supported by the Frederick N. Andrews Fellowship and the SIRG grant administered through the Institute for Cancer Research. R. A. Shelley and ER. D. Chianis were supported by the Institute for Cancer Research summer undergraduate research program funded by the Carroll County Cancer Association. B. L. Allen-Petersen was supported by the NIH NCI 1K22CA237620-01A1 and the Ross-Lynn Scholars Research Grant.

## SUPPLEMENTAL FIGURE LEGENDS

**Supplemental Figure 1.**
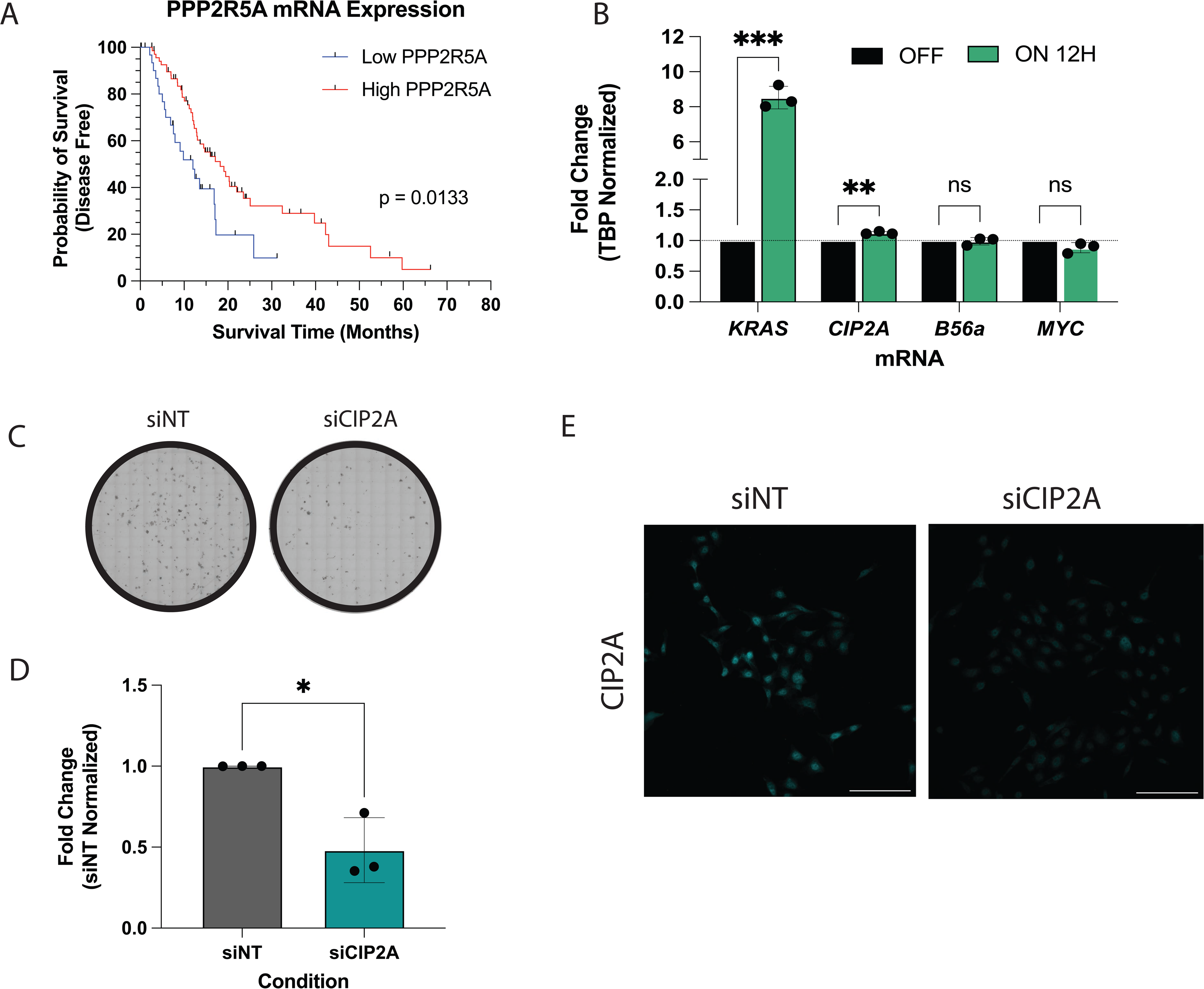
**(A)** Disease-free survival of patients stratified by expression of *PPP2R5A* (B56α) mRNA. Graph generated from publicly available data on CBioPortal TCGA. **(B)** Fold change in mRNA expression of indicated genes in response to 12H of 50ng/mL doxycycline. **(C)** Representative images of clonogenic assay performed with HPDE-iKRAS cells with either siNT or siCIP2A treatment. **(D)** Quantification of three independent replicates of clonogenic assay represented in C. **(E)** Representative image of siCIP2A knockdown in cells used for PLA experiments. Statistical analysis performed using student’s t-test or one-way ANOVA. \**p*<0.05 \*\**p*<0.01 \*\*\**p*<0.001 n.s. = not significant. Scale bar represents 100μm.

**Supplement Figure 2.**
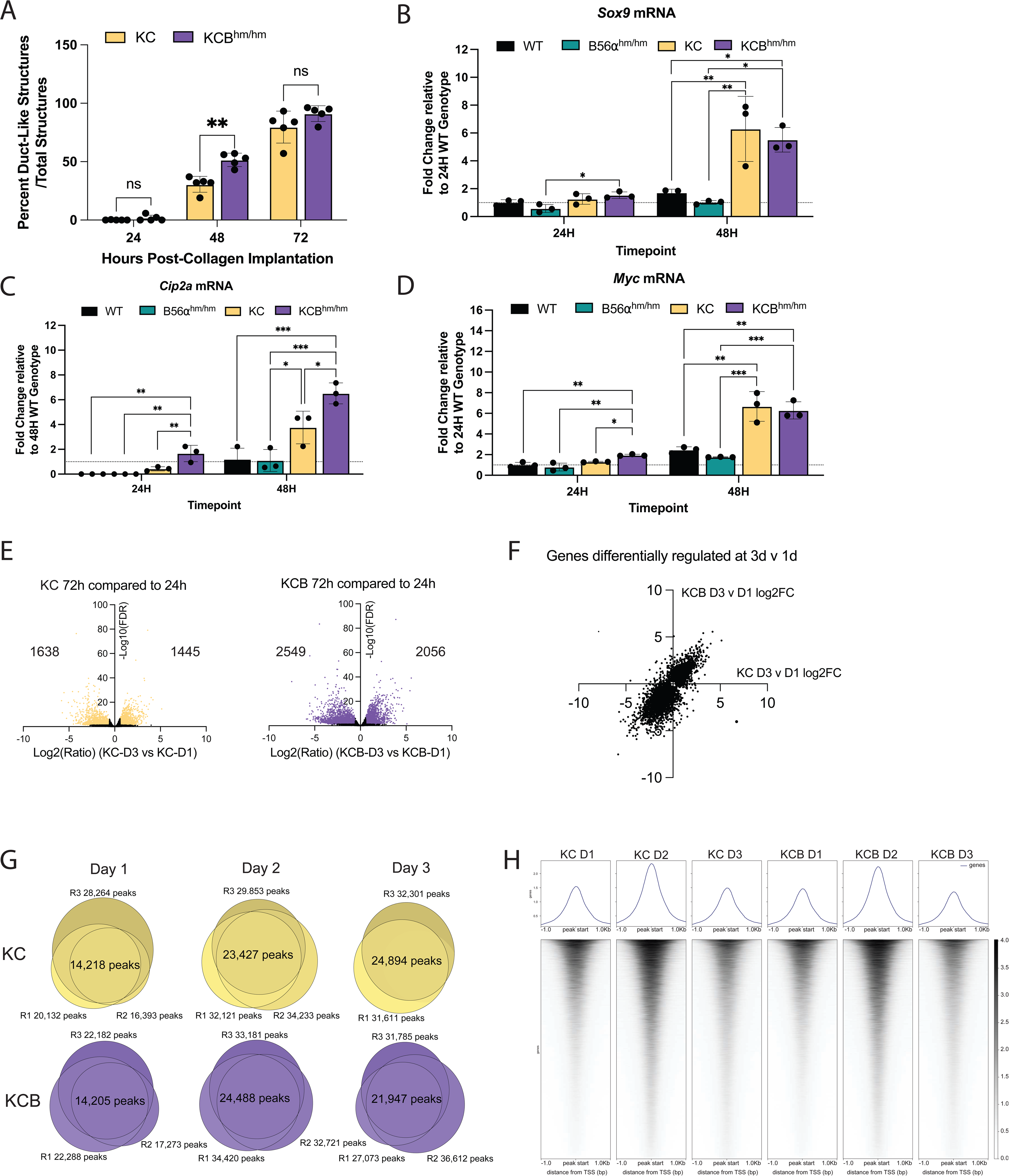
**(A)** Percent of total structures that appear duct-like after 24, 48, and 72H in collagen. Replicates representative of five independent replicates. **(B-D)** mRNA expression of *sox9*, *cip2a,* and *myc* in each genotype at 24, 48H timepoints. **(E)** DEGs identified in KC (left) and KCB (right) at D3 compared to D1. Log2 FC indicates the mean expression level for each gene. Each dot represents one gene. **(F)** Correlation of DEGs regulated in D3 vs D1 between KC (x-axis) and KCB (y-axis). **(G)** Overlapping peaks identified in each replicate of each timepoint and condition for quality control **(H)** Accessible chromatin identified in D1, D2, and D3 of each genotype. Statistical analysis performed using student’s t-test or one-way ANOVA. \**p*<0.05 \*\**p*<0.01 \*\*\**p*<0.001 n.s. = not significant

**Supplemental Figure 3.**
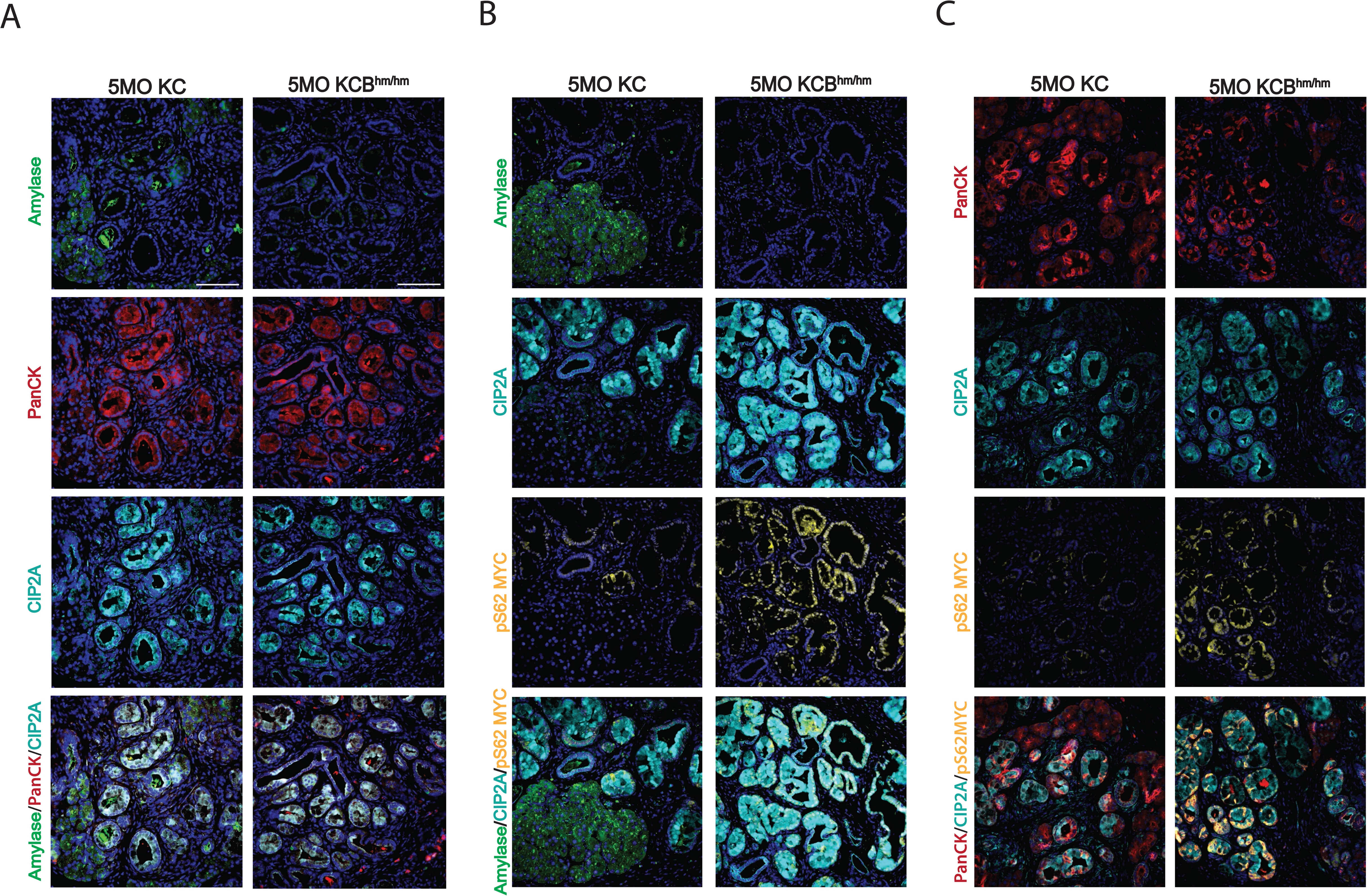
Representative images of pancreata harvested at 5MO from KC or KCB^hm/hm^ mice stained with **(A)** Amylase, PanCK, and CIP2A, **(B)** Amylase, CIP2A, and pS62 MYC, and **(C)** PanCK, CIP2A, and pS62 MYC. All images represent at least three independent biological replicates. Scale bar represents 100μm.

