## Supplementary figures and images for "KRAS-mediated upregulation of CIP2A promotes suppression of PP2A-B56α to initiate pancreatic cancer development"

### Supplemental Figures

Supplemental 1

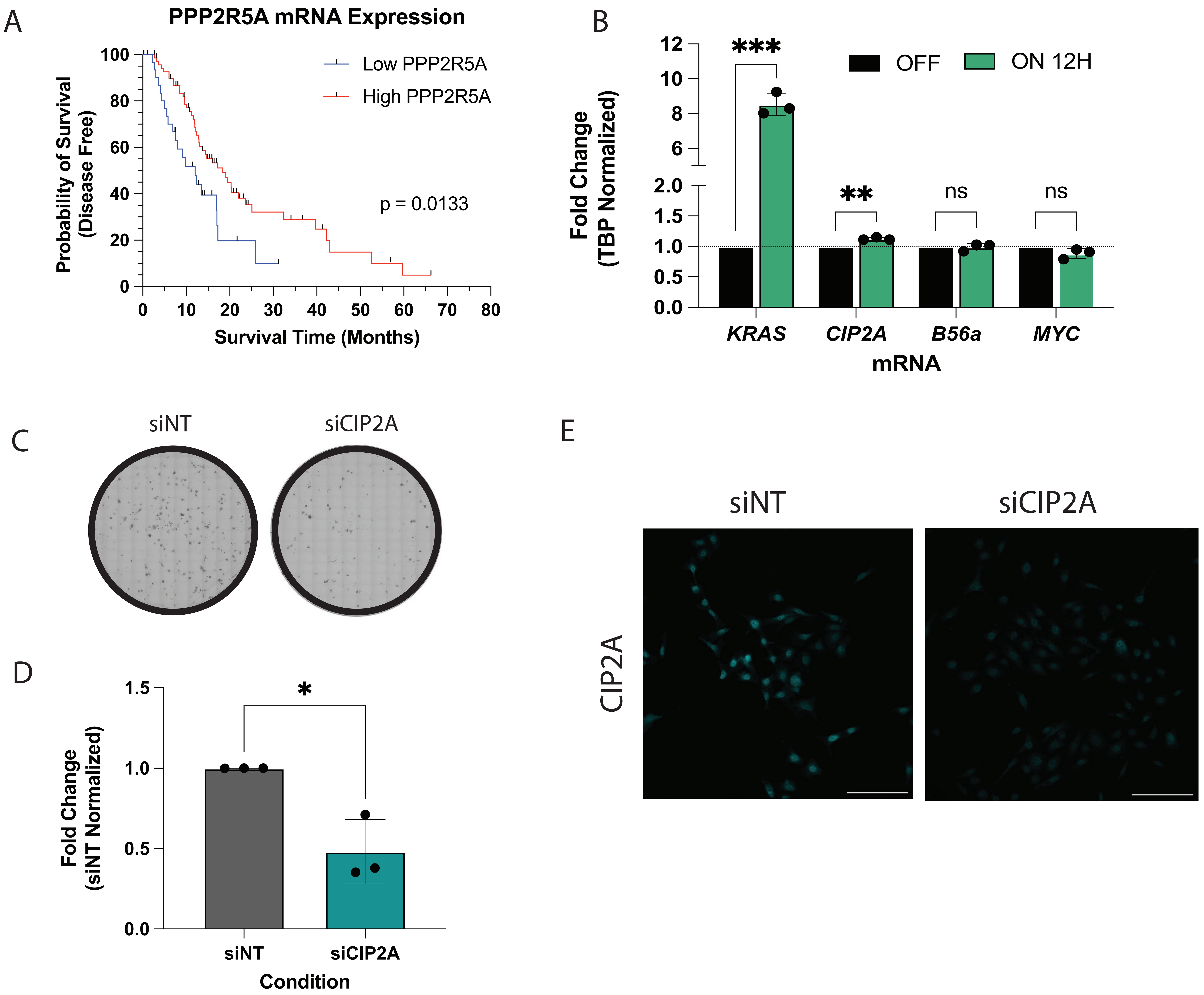

#### Supplemental 2

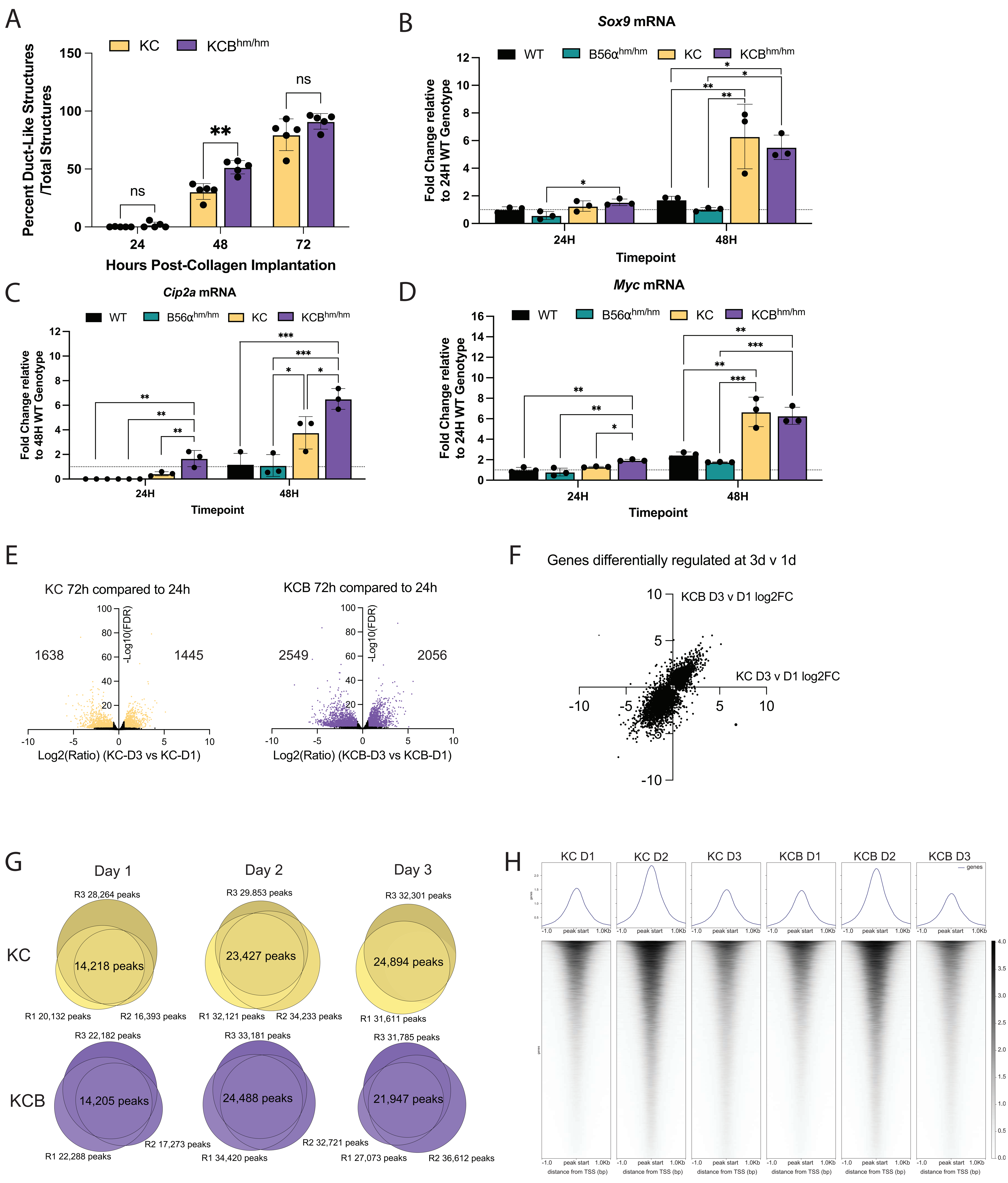

Supplemental 3

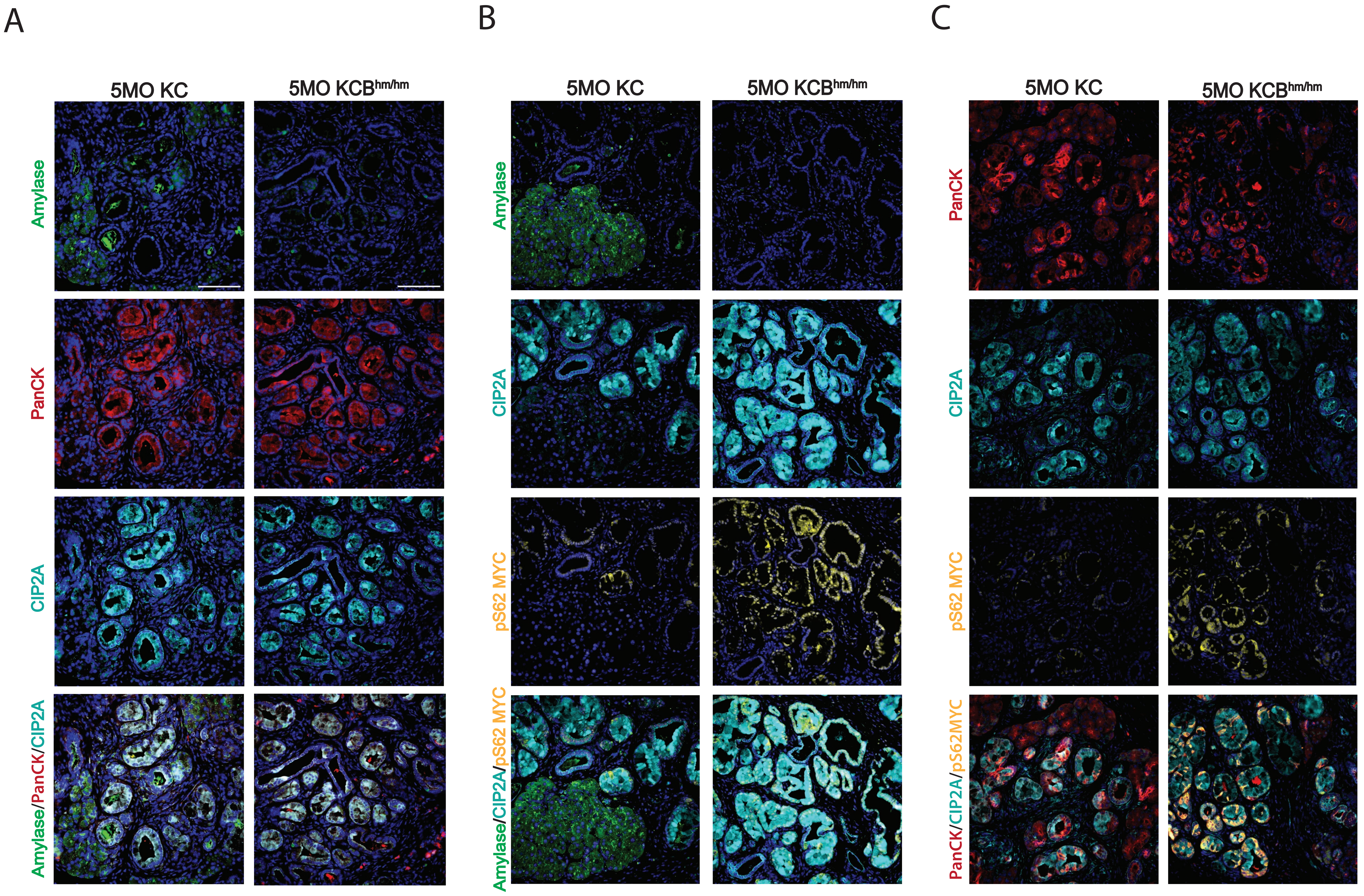
